# Cytidine deaminase protects pancreatic cancer cells from replicative stress and drives resistance to DNA-targeting drugs

**DOI:** 10.1101/2021.10.23.465566

**Authors:** A. Lumeau, N. Bery, A. Francès, M. Gayral, C. Ribeyre, C. Lopez, A. Névot, S. Elkaoutari, G. Labrousse, M. Madrid-Mencia, M. Pillaire, V. Pancaldi, V. Bergoglio, N. Dusetti, J. Hoffmann, L. Buscail, M. Lutzmann, P. Cordelier

**Author notes:** corresponding author. Pierre Cordelier, Cancer Research Centre of Toulouse (CRCT), Team 10: “therapeutic innovation in pancreatic cancer”, 2, avenue Hubert Curien, 31100 Toulouse, France. All authors have seen and approved the manuscript, and that it hasn’t been accepted or published elsewhere.

## Abstract

Chronic DNA replication stress and genome instability are two hallmarks of cancer that fuel oncogenesis and tumor diversity. Therapeutic approaches aimed to leverage tumor-specific replication stress to intolerable levels or to expose vulnerabilities for synthetic lethality purposes have recently gained momentum, especially for pancreatic cancer, a disease with no cure. However, the current knowledge regarding the molecular mechanisms involved in the replication stress response in pancreatic tumors is limited. Cytidine deaminase (CDA) is involved in the pyrimidine salvage pathway for DNA and RNA synthesis. Loss of CDA induces genomic instability in Bloom Syndrome, and CDA protects tumor cells from chemotherapy with pyrimidine analogs. Here, we show that CDA is overexpressed in genetically unstable pancreatic tumors, associates with a DNA replication signature, and is instrumental for experimental tumor growth. In cancer cells, CDA promotes DNA replication, increases replication fork speed, and controls replication stress and genomic stability levels. CDA expression is predictive of DNA-damaging drug efficacy and targeting CDA relieves resistance to chemotherapy in patients models, both *in vitro* and *in vivo*. Our findings shed new light on the mechanisms by which pancreatic cancer cells control replication stress, and highlight targeting of CDA as a potential therapeutic strategy to defeat tumor resistance to treatment.

## INTRODUCTION

At each cell division, billions of nucleotides of DNA must be accurately polymerized, and ensuring that this process occurs prior to the next cell cycle is essential for cellular homeostasis^1^. The DNA replication machinery successfully carries out accurate genome duplication in the face of numerous obstacles, many of which cause DNA replication stress. Replication stress is defined as any hindrance to DNA replication that either stalls, blocks, or terminates DNA polymerization. Due to high levels of proliferation and a deregulated cell cycle, DNA replication stress represents one of the main drivers of clonal diversification, creating the multi-layered genomic instability often seen in cancer^2^. Consequently, cancer cells have evolved to fine-tune DNA replication stress, notably following oncogene activation^3^, for the acquisition of genetic heterogeneity while preserving favorable genotypes that are compatible with tumor growth. This dependence can be exploited for a therapeutic benefit as cancer cells can be pushed towards cell death by increasing DNA damage, without necessarily killing normal cells that have an intrinsically lower DNA damage level and efficient DNA damage repair pathways.

Pancreatic adenocarcinoma (PDAC) is a disease with no cure that will soon rank second in death by cancer worldwide^4^. PDAC is dominated by mutations in *KRAS*, *TP53*, *CDKN2A*, and *SMAD4*, with a handful of gene mutated in 5%–15% of cases, and a number of infrequently mutated genes in most PDAC patients^5,6^. So-called unstable genotypes with numerous structural variations are present in a significant proportion of PDAC patients (10-15%), characterized by the inactivation of DNA maintenance genes (*BRCA1*, *BRCA2*, or *PALB2*) and somatic mutations in genes involved in DNA repair (*BRCA2*, *BRCA1*, *PALB2*, *ATM*, and *RAD51*)^5,6^. This genetic diversity may explain why PDAC has been historically highly recalcitrant to chemotherapies and targeted therapies. However, recent studies have shown that patients with germline mutations in *BRCA2* demonstrate therapeutic response following a first line platinum-containing FOLFIRINOX regimen, followed by PARP inhibitor (Olaparib) treatment^7^. In a recent work, Dreyer et al further classified PDAC experimental models using a DNA repair deficiency (DDR) signature and found that a subset of PDAC samples with squamous transcriptomic profile show evidence of high replication stress, that predicts response to ATR and WEE1 inhibitors^8^. Collectively, these works demonstrate that a significant fraction of PDAC tumours show intrinsic evidence of genetic instability, and that replication stress is beginning to be explored as an attractive target in the clinic^9,10^. However, the molecular mechanisms that govern replication stress in PDAC remain largely unknown.

Cytidine deaminase (CDA) catalyses the irreversible hydrolytic deamination of cytidine and deoxycytidine to uridine and deoxyuridine, respectively, within the pyrimidine salvage pathway for DNA and RNA synthesis (for review^11^). In Bloom syndrome, a genetic disease with one of the strongest known correlations between chromosomal instability and increased risk of malignancy^12^, loss of CDA expression results in a pyrimidine pool imbalance, which in turn generates replication stress leading to segregation defects^18^. In the oncology field, CDA expression is historically recognized as a key chemoresistance factor to cytidine and deoxycytidine analogs such as Ara-C, gemcitabine (dFdC), or decitabine (5-Aza-dC) since it catalyzes their trangtgsformation into inactive metabolites that are excreted by cancer cells. As an example, 80% of the intravenous Ara-C bolus is eliminated in the urine, 90% of which is in its uracil form, thus strongly suggesting direct and active “detoxification” by CDA^13^. In PDAC, patients with high CDA activity are 5-fold more likely to progress following gemcitabine-based therapy^14^, and CDA from the tumor microbiome participates in tumor resistance to treatment^15^. However, the role of CDA *per se* in PDAC oncogenesis has never been investigated so far.

For this study, we aimed to consider the role of CDA on DNA replication in the context of PDAC, independently of its role in resistance to gemcitabine treatment. We report here that CDA is overexpressed in PDAC at diagnosis, and that CDA expression in tumors naives of treatment correlates with worse prognosis. Hence, CDA depletion strongly impairs PDAC experimental proliferation and tumor growth, in the absence of gemcitabine. Gene signature analysis demonstrates that CDA expression positively correlates with DNA replication signature in PDAC and is enriched in genetically unstable tumors. In PDAC cells, CDA promotes DNA synthesis by increasing replication fork stability, CDA is nuclear and locates at the DNA replication fork. In addition, we show that CDA activity is essential to dampens replication stress and genomic instability in these cells. Finally, we show that CDA expression drives resistance to DNA targeting drugs in PDAC experimental models, and that targeting CDA both *in vitro* and *in vivo* sensitizes PDAC tumors to genotoxic treatment. Collectively, we demonstrate here that CDA is involved in the regulation of replicative stress levels in PDAC, a novel dependence that can be exploited for a therapeutic benefit.

## RESULTS

### CDA is overexpressed in PDAC and essential to cell proliferation and tumor growth

CDA expression in tumors has been scarcely investigated to date^11^. Previous reports indicate that CDA is upregulated in human PDAC samples^16^ and in animal models for this disease^17^. To further characterize the expression of CDA in PDAC, we first analyzed surgical specimens from patients with this cancer. Matching, normal adjacent parenchyma was obtained for each patient. Using qRT-PCR, we found that CDA mRNA is significantly overexpressed in PDAC tissue (Fig. 1A, 5.1+/-1.5-fold increase, p<0.001). This finding was comfirmed by *in silico* investigation with the curated TCGA_PAAD dataset^18^ (Extended Figure 1A). In recent years, several studies have demonstrated that molecular data can define subgroups in PDAC with distinct biology^19^. Analysis of PDAC transcriptomes have revealed two main molecular subtypes, with classical tumors on one side showing the highest expression of epithelial and adhesion associated genes, as well as high levels of GATA binding protein 6 (GATA6) mRNA, and basal-like tumors on the other side that are predominantly composed of poorly differentiated tumors, more often CDKN2A or TP53 mutated, and that exhibit the worst outcome^20^. We applied the Purity Independent Subtyping of Tumors (PurIST) classifier^21^ to PDAC tumors from TCGA with known CDA mRNA levels and found that CDA expression is significantly enriched in PDAC primary cells with basal phenotype (Fig. 1B 2.3+/-0.1-fold increase, p<0.001). We extended this analysis to a recently published primary cell lines collection^8^, and found that CDA mRNA is significantly overexpresed in PDAC primary cells with the squamous phenotype, that largely corresponds to the basal-like molecular phenotype (Extended Fig.1B, 1.2+/-0.04-fold increase, p<0.05). PDAC patient tumors from TCGA were next classified as CDA high-expressing (top 25% quartile) or CDA low-expressing (bottom 25% quartile); we found that CDA expression is positively associated with shorter survival (Fig. 1C, p<0.03, hazard ratio = 2.021,). These results indicate that PDAC tumors with the most aggressive molecular phenotype and the shortest survival overexpress CDA.

**Figure 1.**
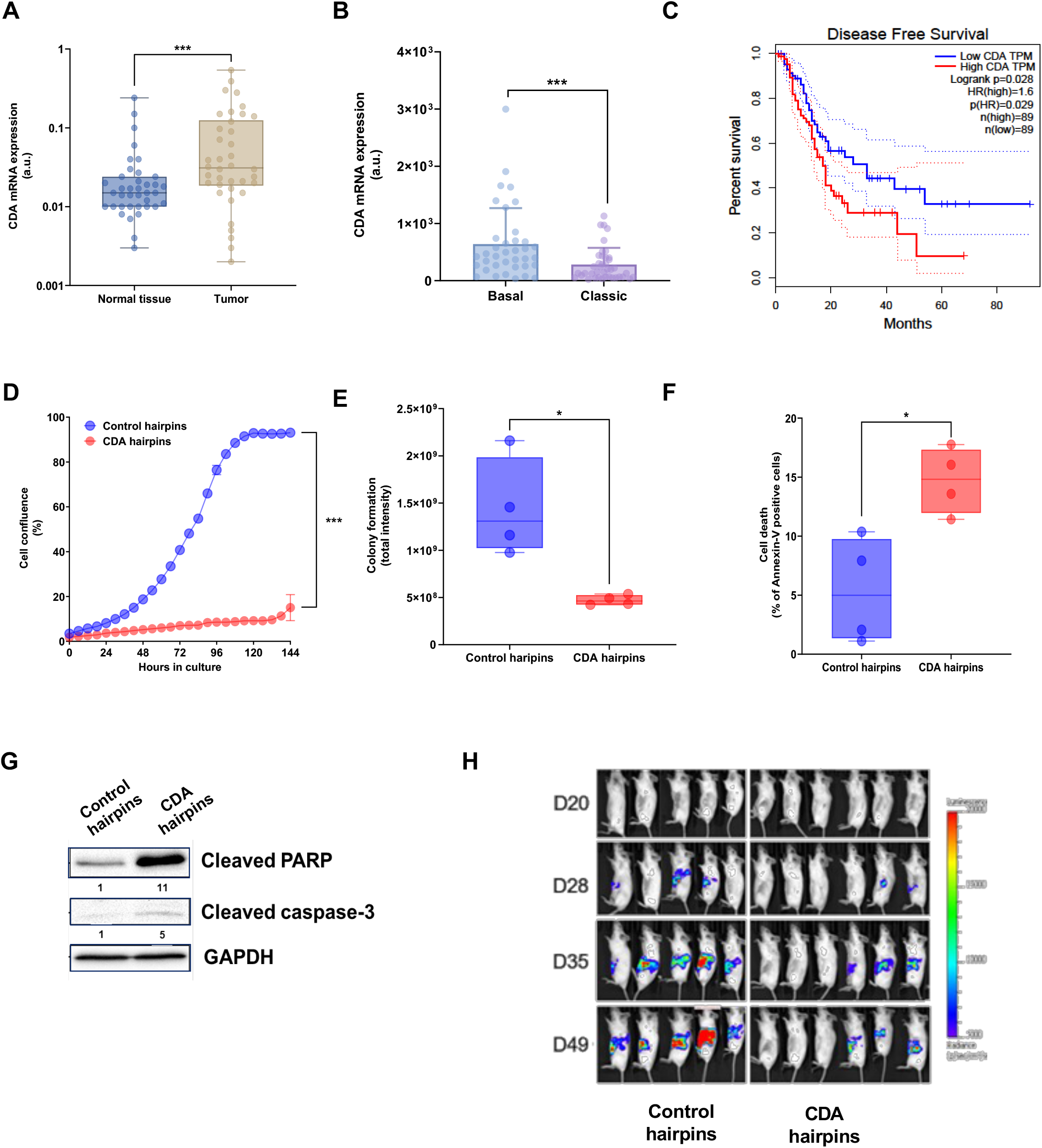
CDA is overexpressed in PDAC tumors and essential to cell proliferation and tumor growth. **A**. Expression of CDA mRNA in pancreatic tumors and matched normal tissue (n=37) from CHU Toulouse and Idibell cohorts. ***p<0.001 (Wilcoxon-Mann-Whitney test). B. CDA mRNA expression in PDAC tumors from TCGA with basal (n=37) or classic (n=40) molecular subtype. p<0.005. **C**. Kaplan Meyer survival plot for CDA expression in PDAC patients (n=41). **D**. Proliferation of CDA-depleted and control MIA PaCa-2 cells measured with the IncuCyte Zoom. Representative of at least three independent experiments performed in triplicate. ***p<0.001 (unpaired t-test) **E**. Colony formation assay of CDA-depleted vs MIA PaCa-2 control cells. Representative of four independent experiments performed from four different transductions and mean quantification of colony areas. *: p-value < 0.05 respectively (unpaired t-test). **F**. Annexin V staining of MIA PaCa-2 pancreatic cancer cells, stably expressing or not CDA hairpins. Mean of four independent experiments from three different transductions. *: p-value < 0.05 (unpaired t-test). **G**. Western blotting of key apoptotic proteins: cleaved PARP, cleaved caspase 3 in CDA-depleted vs MIA PaCa-2 control cells. Representative of seven and five independent experiments respectively. **H**. Representative Tumor formation and growth monitoring in mice with CDA-depleted or control MIA PaCa-2 experimental tumors.

We next addressed the functional importance of CDA in PDAC. We inhibited CDA expression in PDAC cell line using shRNA delivered by lentiviral vectors (Extended Fig. 1C-D) and found that CDA targeting provokes a profound and long-term inhibition of cell proliferation, (−86%±3%, p<0.01, Fig. 1D). We extended this finding to BxPC-3 and Capan-1 PDAC cells (Extended Fig. 1E) or using siRNA pools targeting CDA in MIA PaCa-2 cells (Extended Fig. 1F). In addition, MIA PaCa-2 PDAC cells expressing CDA hairpins showed a reduced ability to form colonies, as compared to control cells (Fig. 1E, −65%±3%, p<0.05). In these cells, CDA invalidation increases cell death by apoptosis 10 foldas monitored by FACS for Annexin-V and Western blotting for caspase-3 and PARP cleavage (Fig. 1F-G). We next engrafted human PDAC cell expressing CDA hairpins in the pancreas of athymic mice. Mice receiving PDAC cells expressing control hairpins were used as control. We found that silencing CDA in PDAC cells results in 50% mice not developing experimental tumors (Fig. 1H). When tumors developed, CDA silencing significantly decreased tumor growth, as compared to control tumors (−97%±17%, p<0.01, Fig. 1H and Extended Figure 1G). Collectively, these data indicate that PDAC cell proliferation and tumor growth strongly rely on CDA expression.

### CDA increases replication fork speed and restart efficiency in PDAC cells

To gain further insights into the role of CDA in PDAC growth, we performed gene signature enrichment analysis in the quartile of PDAC tumors from TCGA overexpressing this enzyme. As shown in Fig. 2A and Table 1, CDA expression positively correlates with the transcriptomic signature of DNA replication (normalized enrichment score = 2.28, p<0.01). We next performed lentiviral transduction of CDA in MIA PaCa-2 PDAC cells and verified efficient mRNA (Extended Fig.2 A) and protein (Extended Fig.2B) expression and activity (Extended Fig. 2C). Cells expressing luciferase were used as control. We found that CDA expression induces the DNA replication signature in these cells (Fig. 2B, normalized enrichment score = 1.98, p<0.01 and Table 2). We next addressed whether CDA could act on DNA replication in PDAC cells. We performed DNA spreading analysis to visualize and follow the spatial and temporal progress of individual DNA replication forks. The method relies on the detection of incorporated thymidine analogues during DNA synthesis in the S phase of the cell cycle by indirect immunofluorescence following cell lysis and DNA fibers spread (Fig. 2C). Thus, PDAC cells expressing CDA were incubated with IdU and CldU thymidine analogs. Cells expressing luciferase were used as control. Figure 2D demonstrates that DNA tracks are significantly longer in cells overexpressing CDA as compared to control cells (12.5μm±0.37 μm *vs* 9.4μm±0.34μm, p<0.001), indicating that replication forks speed is increased in CDA-expressing PDAC cells.

**Figure 2.**
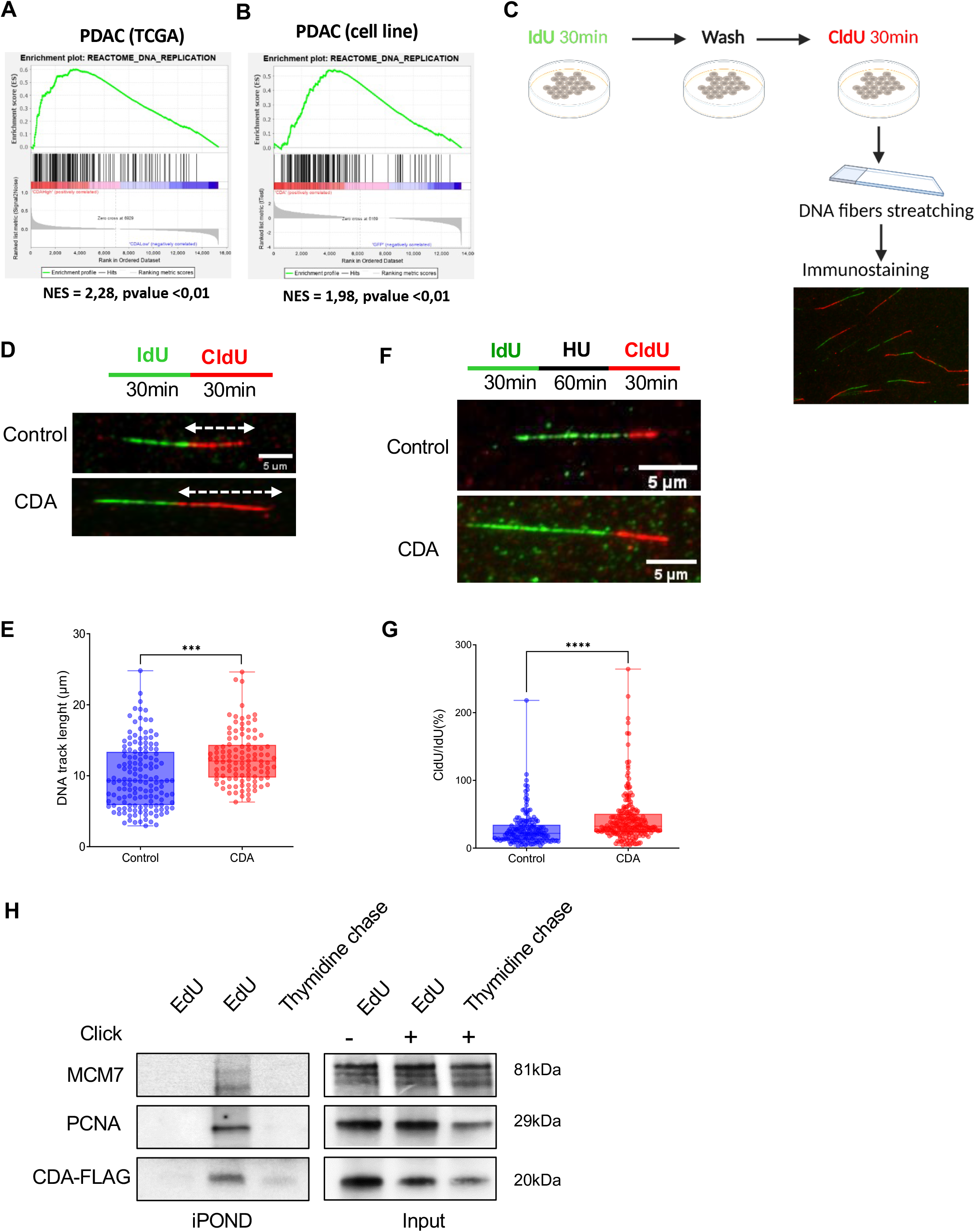
CDA increases replication fork speed and restart efficiency in PDAC cells. Enrichment plots for DNA replication pathway (GSEA-Reactome) of (**A**) CDA high expressing TCGA PDAC samples (high CDA n=39, low CDA n=38) and (**B**) MIA PaCa-2 cells overexpressing CDA. Such enrichment was not found in cells overexpressing the catalytic mutant of CDA. Representative of 5 independent transduction pools. **C.** Schematic procedure of DNA stretching experiment. IdU = 5-Iodo-2’-deoxyuridine, CldU = 5-chloro-2’-deoxyuridine. **D.** Immunofluorescence detection of IdU (green) and CldU (red) DNA tracks after DNA stretching of MIA PaCa-2 overexpressing or not CDA, white arrows show the measured red tracks. **E**. Quantification of the length of DNA tracks (μm), at least 100 DNA fibres were measured per condition. ***: p<0.001 (Wilcoxon-Mann-Whitney test). **F.** Immunofluorescence detection of IdU (green) and CldU (red) DNA tracks after DNA stretching of control MIA PaCa-2 cells, or cells overexpressing CDA, white arrows show the measured CldU tracks. HU: Hydroxyurea, treatment at 4mM for 1h. **G**. Quantification of the length of CldU tracks after HU treatment (μm), at least 70 DNA fibres were measured per condition. Results are representative of 3 independent transduction pools. ****: p<0.0001 (Wilcoxon-Mann-Whitney test). **H**. IPOND experiment in MIA PaCa-2 cells overexpressing CDA-FLAG. PCNA and MCM7 were used as control. Cells were treated with EdU and revealed or not with Click-it EdU (EdU- and EdU+, respectively). After EdU incorporation, cells were treated with Thymidine to chase EdU and to address whether candidate proteins progress with replication fork. Results are representative of four independent experiments.

**Table 1.**
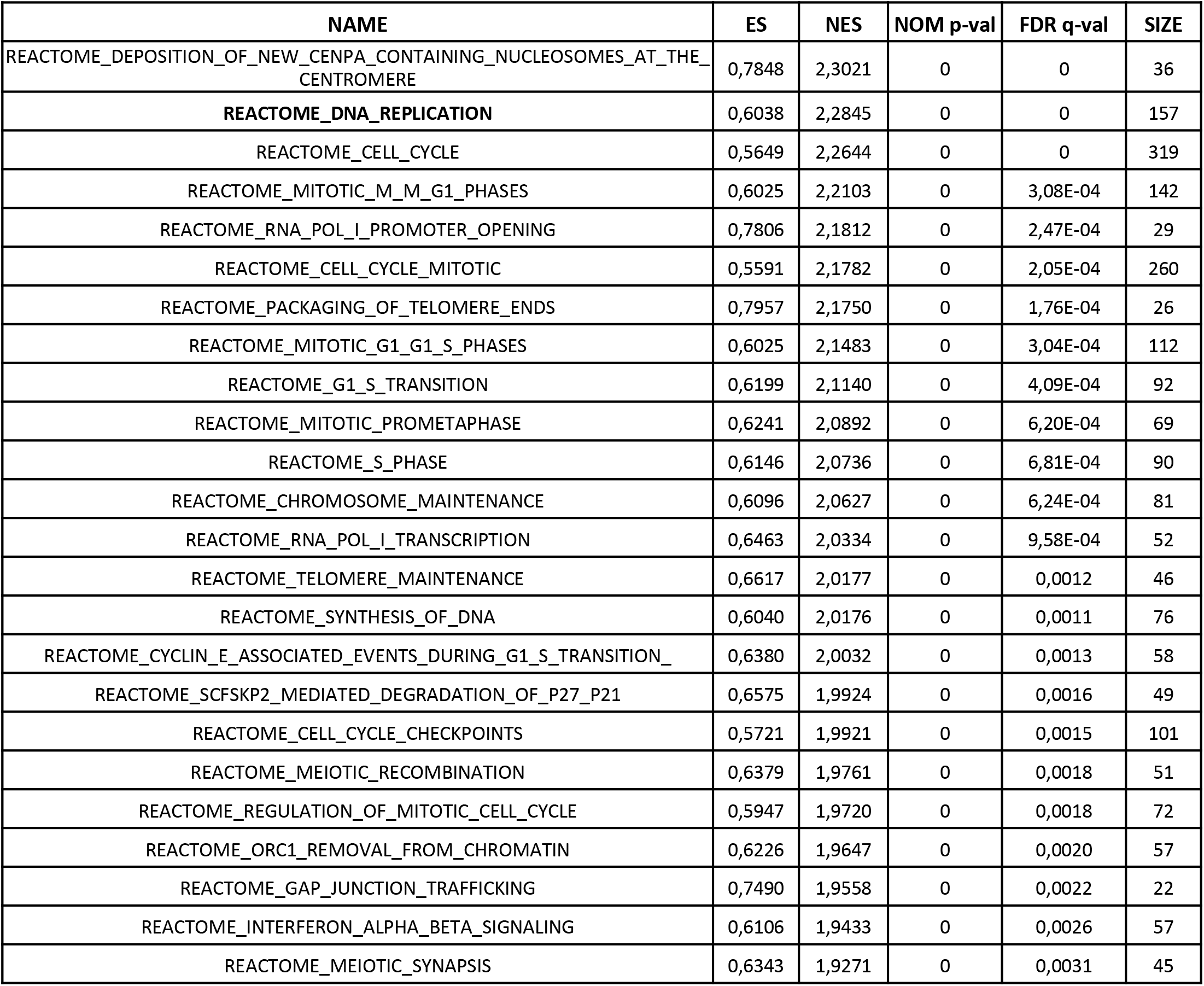
25 most enriched pathways in CDA high vs CDA low TCGA PDAC samples. ES: enrichment score, NES: normalized enriched score. Gene Set Enrichment Analysis using Reactome gene set (n=39 high CDA, n=38 low CDA).

**Table 2.**
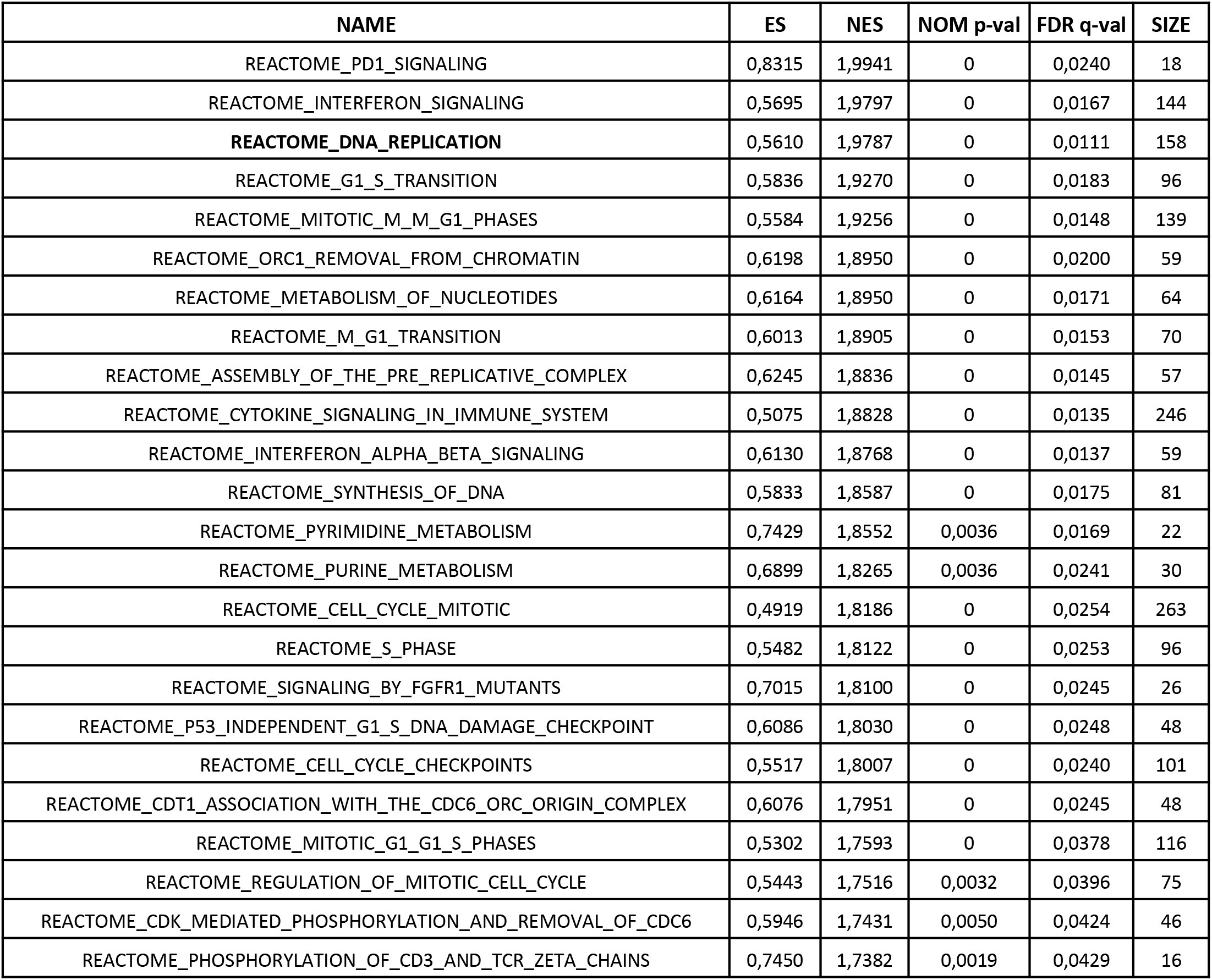
25 most enriched pathways in CDA vs CTRL overexpressing MIA PaCa-2. ES: enrichment score, NES: normalized enriched score. Gene Set Enrichment Analysis using Reactome gene set, results obtained from five independent transduction pools.

We next investigated DNA replication fork restart in PDAC cells expressing CDA. To induce fork stalling, we used hydroxyurea (HU) that inhibits riboside nucleotide reductase (RNR) and causes the depletion of deoxynucleotide triphosphates (dNTPs). PDAC cells were incubated with thymidine analog IdU, to measure initial fork speed, followed by treatment with HU and pulse with CldU. Here, the length of the CldU tract after HU treatment served as a measure of fork restart efficiency. We found that cells expressing CDA show significantly more CldU positive DNA fibers, as compared to control cells, indicative of increased replication fork restart following HU treatment (Fig. 2F, Extended Fig. 2D, 91.7%±1.7% *vs* 78.7%±1.3%, respectively, p<0.001). In addition, the increase in CldU/IdU ratio indicates that CDA increases replication fork speed following stalling (Fig. 2F-G, 43.3%±2.5% *vs* 28.2%±1.9%, respectively, p<0.001).

We next addressed whether CDA could physically interact with the DNA replication forks machinery to increase DNA synthesis. Following cell fractionation of cells expressing FLAG-tagged CDA, we demonstrate that CDA locates both in the cytoplasm, but also in the nucleus of cells overexpressing the enzyme (Extended Fig.2E). Then, we performed Isolation of Proteins On Nascent DNA (iPOND) analysis. Briefly, cells are incubated with the thymidine analogue EdU to label newly replicated DNA. After cross-linking of proteins and DNA with formaldehyde, the click reaction is performed to link biotin to EdU. After cell lysis and sonication to shear chromatin, proteins in proximity to biotin and EdU-labeled DNA are purified with streptavidin-coated agarose beads and resolved by Western blotting (Extended Fig. 2F). Results presented in Fig. 2H show that MCM7 and PCNA, two major components of the fork replisome, are detected within the EdU positive fraction, indicating interaction with newly synthesized DNA. Remarkably, we also located CDA at the DNA replication fork (Fig. 2F, bottom panel). We confirmed this finding in Hela cells that express relatively high levels of CDA (Extended Fig. 2G). We explored another setting in the iPOND experiment and replaced EdU pulse with a chase of thymidine to track how proteins assemble and disassemble from a nascent DNA segment (Fig. 2F, right). In this condition, both MCM7, PCNA and CDA signal disappears, indicating that these proteins progress with the DNA replication machinery. The latter result indicates that CDA is progressing with the replisome rather than just being a protein constitutively bound on chromatin. Collectively, we provide herein the first demonstration that CDA stimulates DNA synthesis, increases replication fork restart efficiency, and is located at the DNA replication fork in PDAC cells.

### CDA controls replication stress levels in PDAC cells

CDA loss in Bloom Syndrome cells participates in the formation of replication defects and sister chromatid exchange^22^. Considering the newly described role of CDA on DNA replication, we investigated whether CDA may participate in DNA replication stress control in PDAC cells. We measured DNA replication of cells expressing or not shRNA against CDA. Internal control consisted in cells expressing control hairpins and treated with the DNA polymerase inhibitor aphidicolin. As shown in Fig. 3A, both CDA targeting and aphidicolin treatment significantly decrease the length of DNA tracks (Fig. 3A-B, −39% and −62%, p<0.001, respectively), highlighting a reduction of replication fork speed. Transcriptomic analysis indicates that the ATR response to replicative stress signature is enriched in response to CDA targeting in PDAC cells, when compared to control cells (Fig. 3B, normalized enrichment score = 1.76, p<0.01, Table 3). This finding is further supported by the strong activation of the Chk1 effector kinase in MIA PaCa-2 cells (Fig. 3C), Capan-1 (Extended Fig. 3A) and BxPC-3 (Extended Fig. 3B) cells expressing hairpins against CDA, as compared to control cells. We next measured γ-H2AX foci in S-phase cells as a canonical marker of DNA breaks and indicative of replication stress. As indicated in Fig. 3E and 3F, CDA expression in MIA PaCa-2 cells significantly decreases γ-H2AX foci in EdU-positive PDAC cells, as compared to control cells (Fig. 3F, −48%±7, p<0.001). This effect is entirely dependent on CDA deaminase activity, as the expression of a catalytically inactive mutant of the enzyme has no impact on γ-H2AX staining (Fig. 3E-F). On the contrary, silencing CDA in BxPC-3 cells results in γ-H2AX accumulation in S-phase cells (Fig. 3G, 2+/-0.02-fold increase, p<0.001). We extended this finding to Capan-1 cells expressing hairpins against CDA (Extended Fig. 3C), and to MIA PaCa-2 cells treated with pharmacological inhibitors of CDA (Extended Fig. 3D) or incubated with cytidine and/or deoxycytidine to phenocopy pyrimidine pool imbalance following CDA deficiency (Extended Fig. 3E). RPA (replication protein A) protects exposed single-stranded DNA during DNA replication, accumulates in response to replicative stress, and is phosphorylated at single-strand breaks. Remarkably, we found that CDA expression decreases the number of RPA foci and p-RPA intensity in PDAC cells (Extended Fig 3F, −67%±8%, p<0.001,).

**Figure 3.**
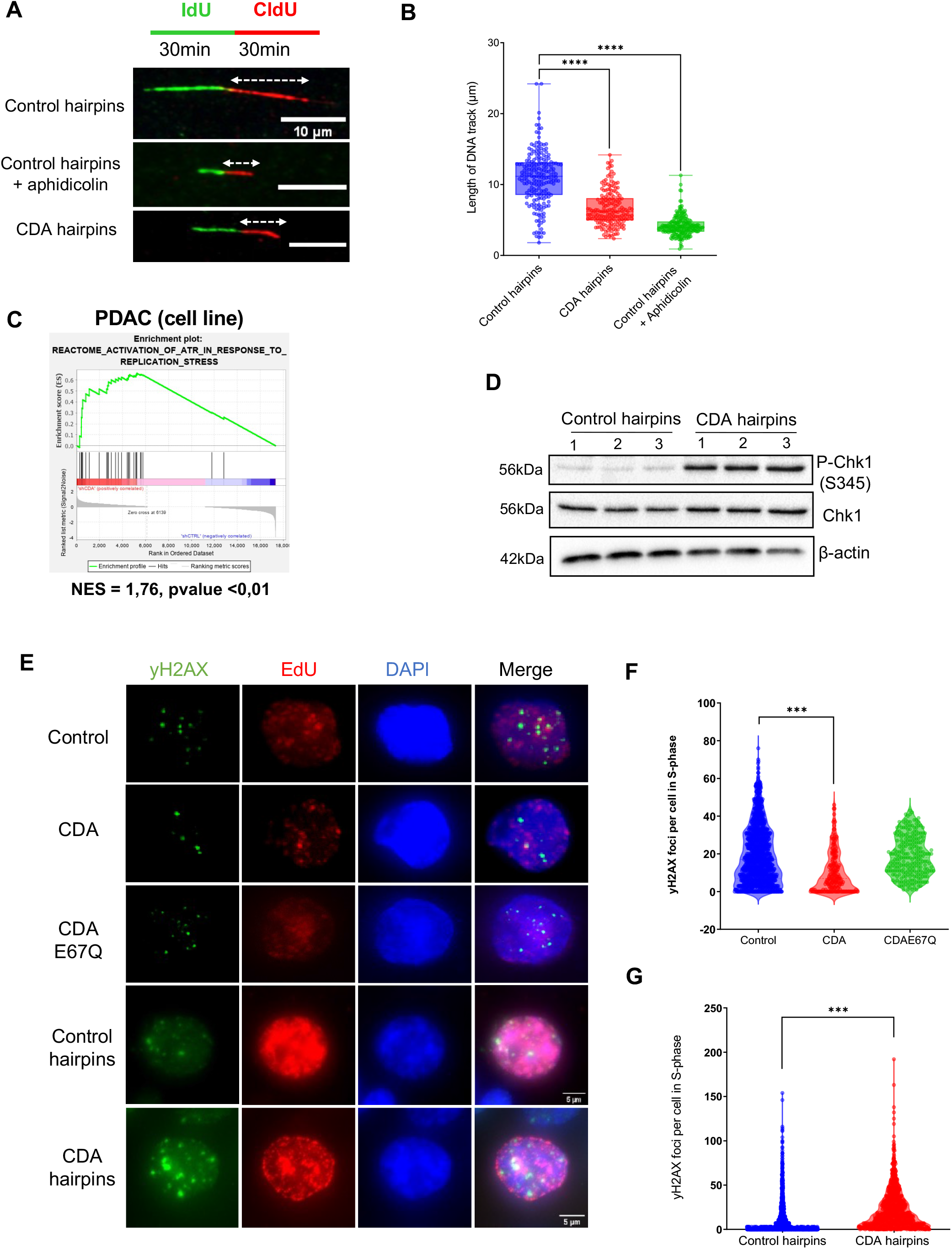
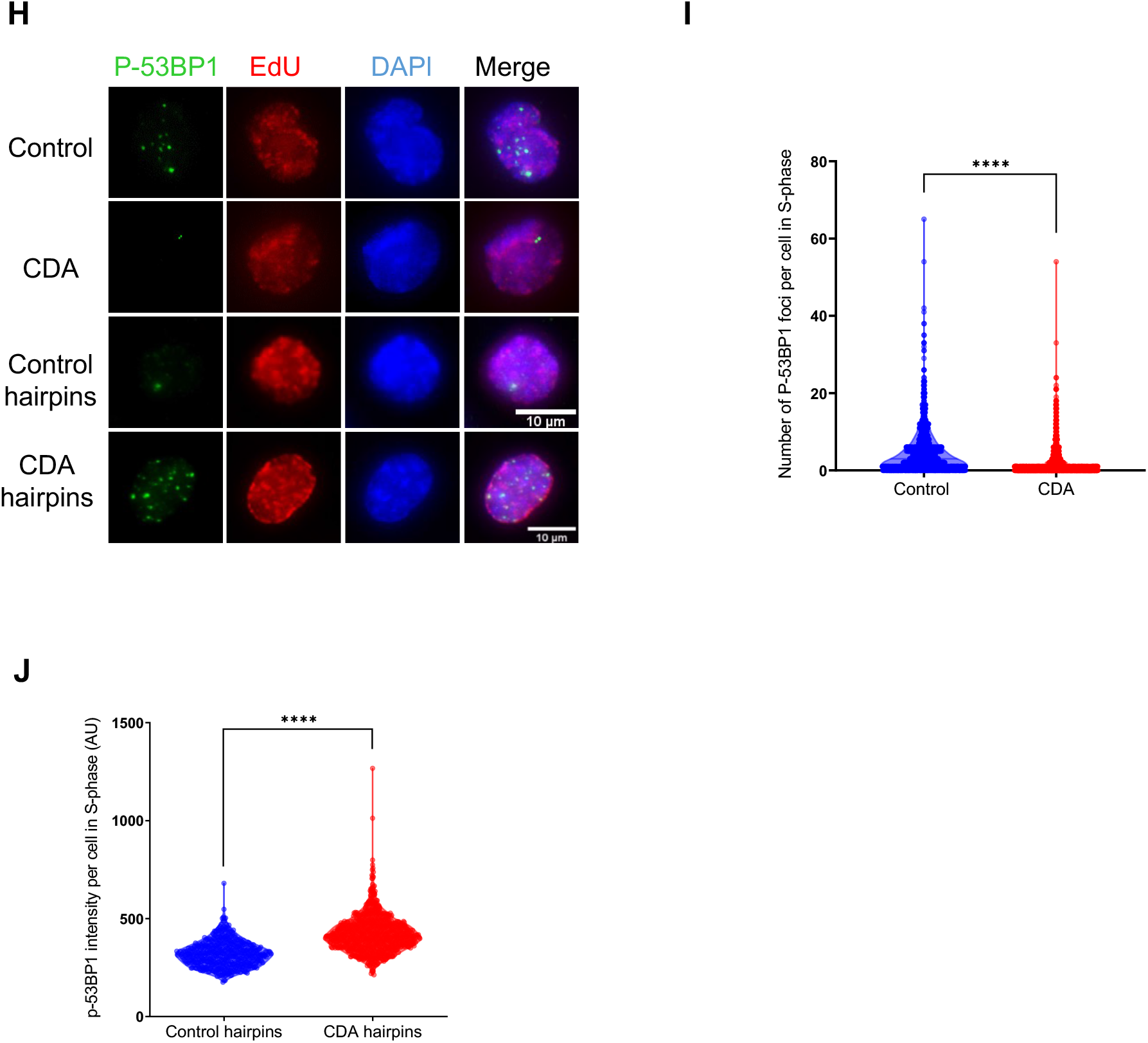
CDA controls replication stress levels in PDAC cells. **A.** Immunofluorescence detection of IdU (green) and CldU (red) DNA tracks after DNA stretching of MIA PaCa-2 cells expressing or not CDA hairpins, white arrows show the measured red tracks, Aphidicoline treatment 24h at 0,2μM was used as a positive control of DNA elongation inhibition, IdU: iodo deoxyuridine, CldU: chloro-deoxyuridine. **B**. Quantification of the length of DNA tracks (μm), at least 150 fibres were measured per condition. Results representative of three independent experiments. ****: p<0.0001 (Mann & Whitney test). **C.** Enrichment plot for Reactome activation of ATR in response to replication stress pathway in MIA PaCa-2 cells depleted for CDA (CDA hairpins), NES = 1,76, p <0.01. Results are representative of three independent transduction pools. **D.** Western Blotting for P-Chk1 (S345), and Chk1 in MIA PaCa-2 cells depleted for CDA or expressing control hairpins. β-actin is used as loading control. Results are representative of three independent pools of transductio*n*. **E.** Immunofluorescence detection of P-H2AX foci (S139) (green), DAPI (blue) and EdU (red). **F**. quantification of yH2AX foci in S-phase cells (EdU+), in at least 500 control cells, of cells overexpressing CDA (MIA PaCa-2) and CDAE67Q (MIA PaCa-2) or depleted for CDA (**G**. CDA hairpins, BxPC-3). Results are representative of three independent transduction pools. ***: p<0.001 (Mann & Whitney test). **H.** Immunofluorescence detection of P-53BP1 (green) and EdU (red). Quantification of the number of p-53BP1 foci and intensity per cell in S-phase (EdU+, red) in at least 500 cells control cells, in MIA PaCa-2 overexpressing CDA (CDA, **I**) or BxPC-3 depleted for CDA (CDA hairpins, **J**). Results are representative of three independent transduction pools of MiA PaCa-2 and two independent transduction pools of BxPC-3. ****: p<0.0001 (Wilcoxon-Mann-Whitney test).

**Table 3.** 25 most enriched pathways in CDA hairpins vs CTRL hairpins MIA PaCa-2. ES: enrichment score, NES: normalized enriched score. Gene Set Enrichment Analysis using Reactome gene set, results obtained from three independent transduction pools.

| NAME | ES | NES | NOM p-val | FDR q-val | SIZE |
| --- | --- | --- | --- | --- | --- |
| REACTOME_BETA_DEFENSINS | 0,7506 | 2,1023 | 0,0000 | 0,0012 | 35 |
| REACTOME_HOMOLOGOUS_DNA_PAIRING_AND_STRAND_EXCHANGE | 0,6858 | 1,9156 | 0,0000 | 0,0345 | 34 |
| **REACTOME_ACTIVATION_OF_ATR_IN_RESPONSE_TO_REPLICATION_STRESS** | 0,6609 | 1,7577 | 0,0000 | 0,2422 | 29 |
| REACTOME_G2_M_DNA_DAMAGE_CHECKPOINT | 0,5754 | 1,7292 | 0,0000 | 0,2450 | 49 |
| REACTOME_ACTIVATION_OF_THE_PRE_REPLICATIVE_COMPLEX | 0,6516 | 1,6981 | 0,0102 | 0,2891 | 25 |
| REACTOME_HDR_THROUGH_HOMOLOGOUS_RECOMBINATION_HRR_ | 0,5503 | 1,6852 | 0,0085 | 0,2800 | 54 |
| REACTOME_HDR_THROUGH_SINGLE_STRAND_ANNEALING_SSA_ | 0,6035 | 1,6719 | 0,0027 | 0,2787 | 31 |
| REACTOME_G2_M_CHECKPOINTS | 0,4825 | 1,6363 | 0,0000 | 0,3443 | 113 |
| REACTOME_TP53_REGULATES_TRANSCRIPTION_OF_ADDITIONAL_CELL_CYCLE_GENES_ WHOSE_EXACT_ROLE_IN_THE_P53_PATHWAY_REMAIN_UNCERTAIN | 0,6613 | 1,5787 | 0,0332 | 0,5097 | 18 |
| REACTOME_DIGESTION | 0,6116 | 1,5698 | 0,0196 | 0,4975 | 23 |
| REACTOME_REPRODUCTION | 0,4868 | 1,5602 | 0,0033 | 0,4909 | 73 |
| REACTOME_INTRINSIC_PATHWAY_FOR_APOPTOSIS | 0,5212 | 1,5458 | 0,0112 | 0,5084 | 50 |
| REACTOME_DNA_DOUBLE_STRAND_BREAK_REPAIR | 0,4445 | 1,5454 | 0,0000 | 0,4707 | 111 |
| REACTOME_DNA_REPLICATION_PRE_INITIATION | 0,4764 | 1,5437 | 0,0000 | 0,4439 | 75 |
| REACTOME_MEIOSIS | 0,5073 | 1,5405 | 0,0151 | 0,4257 | 53 |
| REACTOME_DEADENYLATION_OF_MRNA | 0,6215 | 1,5332 | 0,0352 | 0,4225 | 19 |
| REACTOME_ESTROGEN_DEPENDENT_NUCLEAR_EVENTS_DOWNSTREAM_OF_ESR_ MEMBRANE_SIGNALING | 0,6132 | 1,5292 | 0,0230 | 0,4106 | 22 |
| REACTOME_HOMOLOGY_DIRECTED_REPAIR | 0,4639 | 1,5225 | 0,0076 | 0,4098 | 86 |
| REACTOME_FANCONI_ANEMIA_PATHWAY | 0,5616 | 1,5041 | 0,0230 | 0,4504 | 29 |
| REACTOME_PROCESSING_OF_DNA_DOUBLE_STRAND_BREAK_ENDS | 0,4915 | 1,5002 | 0,0058 | 0,4399 | 53 |
| REACTOME_TRANSCRIPTIONAL_REGULATION_BY_E2F6 | 0,5435 | 1,4780 | 0,0320 | 0,4907 | 34 |
| REACTOME_MEIOTIC_RECOMBINATION | 0,5697 | 1,4653 | 0,0455 | 0,5140 | 25 |
| REACTOME_DIGESTION_AND_ABSORPTION | 0,5603 | 1,4616 | 0,0260 | 0,5058 | 28 |
| REACTOME_GAP_FILLING_DNA_REPAIR_SYNTHESIS_AND_LIGATION_IN_GG_NER | 0,5556 | 1,4491 | 0,0469 | 0,5290 | 25 |

The p53-binding protein 1 (53BP1) is a well-known DNA damage response (DDR) factor, which is recruited to nuclear structures at the site of DNA damage notably following DNA replication stress. In MIA PaCa-2 cells overexpressing CDA, we found that phospho-53BP1 (P-53BP1) foci are reduced as compared to control cells (Fig. 3H-I, −56%%±5%, p<0.001,). On the contrary, BxPC-3 cells that express CDA hairpins demonstrate higher levels of P-53BP1 foci in S-phase than control cells (Fig. 3J +30%%±1%, p<0.001). The latter finding not only confirms that CDA depletion provokes DNA replication stress, but also indicates that the DDR pathway is unaltered in these cells. Collectively, these results indicate that replication stress level in PDAC cells depends on CDA expression level.

### CDA controls of genomic instability in PDAC cells

Residual and unsolved replication stress can lead to genetic instability, including in Bloom syndrome cells depleted for CDA^22^. Common fragile sites (CFSs) are large chromosomal regions that exhibit breakage on mitotic chromosomes upon replication stress. They become preferentially unstable at the early stage of cancer development and are hotspots for chromosomal rearrangements in cancers^23^. Recent studies have shown that FANCD2 facilitates replication across CFSs and can be used as a marker of the presence of unstable CSFs in cells subjected to replication stress^24^. Hence, if under-replicated DNA at CFSs persists into late mitosis, it will lead to the formation of ultrafine anaphase bridges (UFBs), causing chromosome non-disjunction and mitotic catastrophes^25^. FANCD2 forms symmetric foci at each end of UFBs and has a role in the resolution of UFBs in mitosis. As shown in Figure 4A-B, MIA PaCa-2 cells highly expressing CDA show significantly less FANCD2 foci in early mitosis (−56%%±14%, p<0.001), while CDA targeting strongly increases the number of leased CFSs in these cells (Fig. 4A-B, +76%%±7%, p<0.001). These results strongly suggest that CDA limits the number of UFBs in PDAC cells because of reduced replication stress.

**Figure 4.**
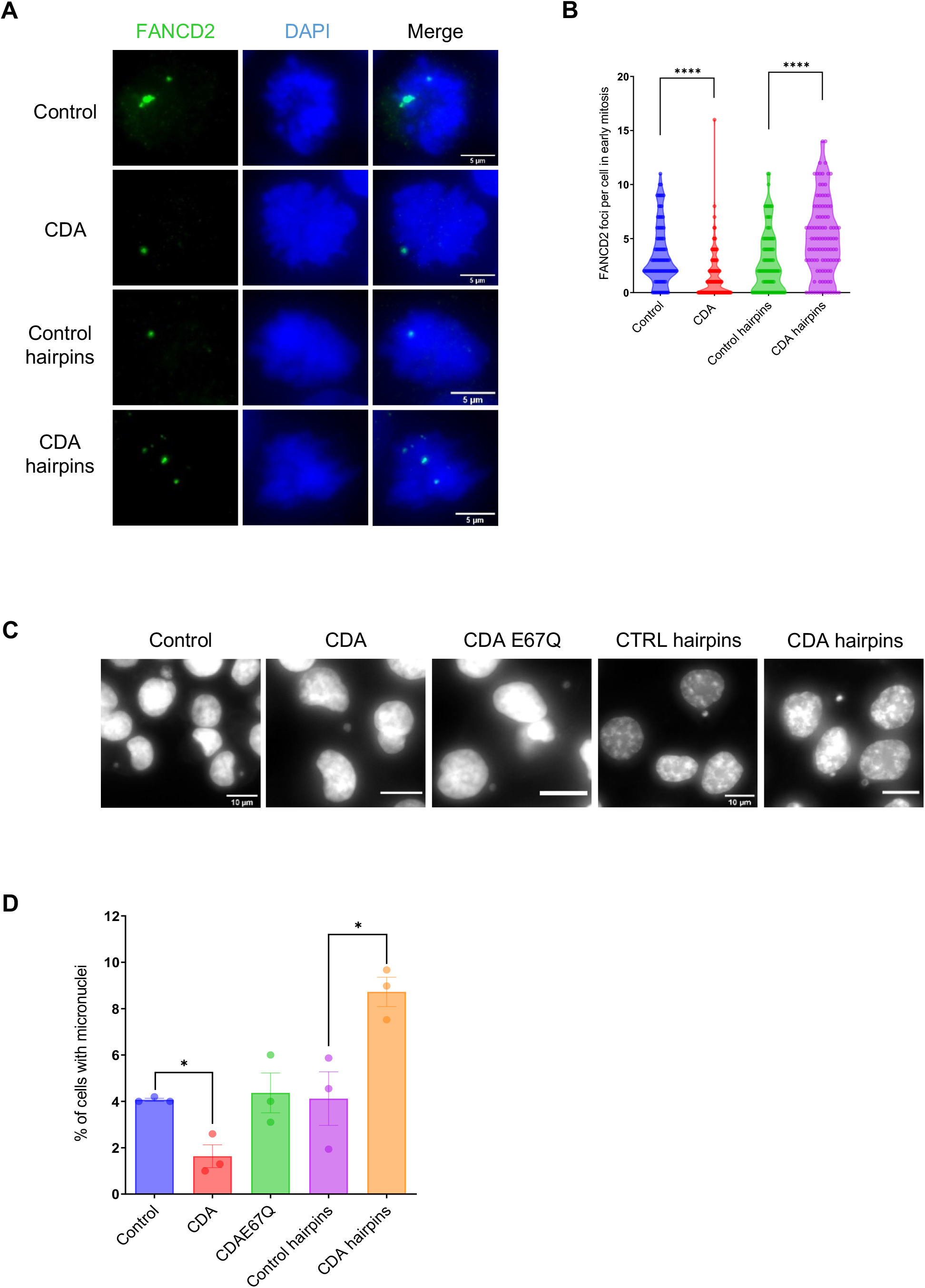
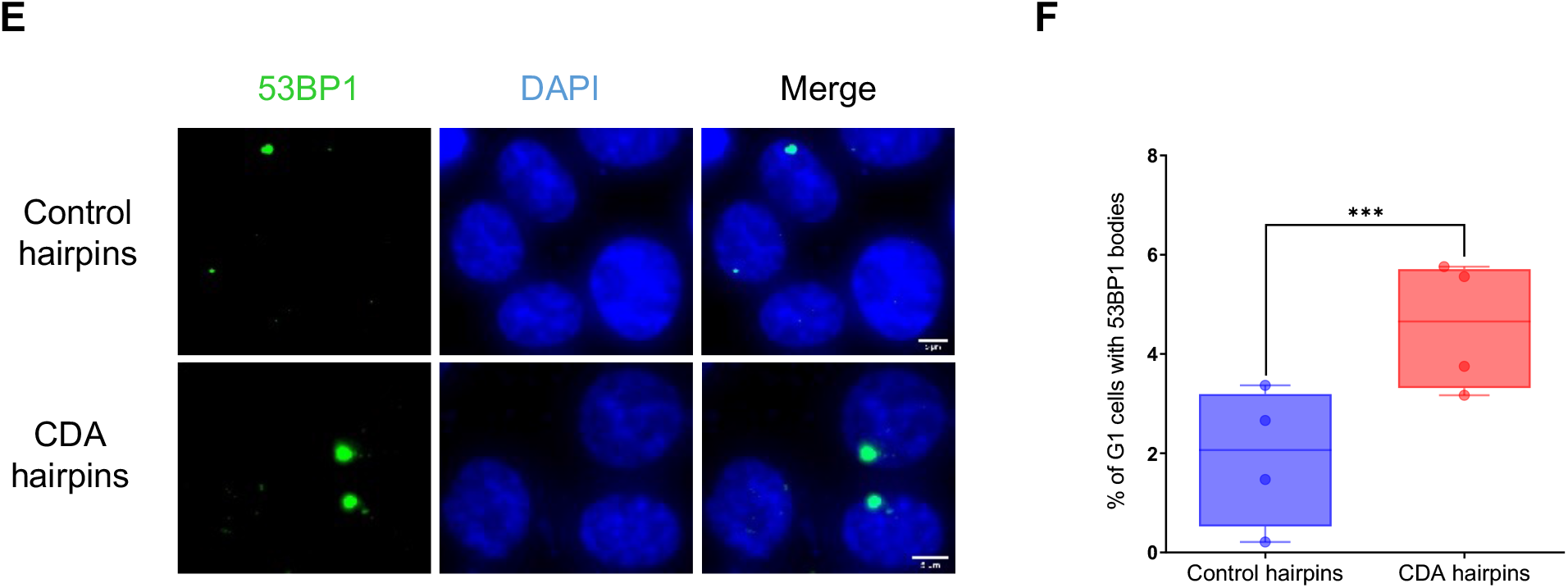
CDA controls genomic instability of PDAC cells. **A.** Immunofluorescence detection of FANCD2 foci (green), DAPI (blue). **B**. Quantification of the number of FANCD2 foci in early mitotic cells *(prophase, metaphase)* in at least 100 control cells, or cells overexpressing CDA (MIA PaCa-2) or depleted for CDA (CDA hairpins, Capan-1). ****: p<0.0001 (Wilcoxon-Mann-Whitney test). **C.** Fluorescence detection of micronuclei using DAPI staining, white arrows show micronuclei. **D**. quantification of the percentage of MIA PaCa-2 and BxPC-3 cells with at least one micronucleus. Results are representative of three independent transduction pools. *: p<0.05 (Wilcoxon-Mann-Whitney test). **E.** Immunofluorescence detection of 53BP1 bodies (green) and DAPI (blue). **F**. Quantification of the number of 53BP1 bodies in G1 cells in at least 1000 control cells and cells expressing CDA hairpins. Results are representative of three independent transduction pools. ***: p<0.001 (Wilcoxon-Mann-Whitney test).

Micronuclei are the small sized nuclei that form from one or a few chromosomes or chromatin fragments that are not incorporated into the daughter nuclei during cell division^26^. Micronuclei formation usually serves as an index of genotoxic effects and chromosomal instability. By microscopy, we found that CDA expression significantly limits the number of DAPI-positive micronuclei in MIA PaCa-2 cells (Fig. 4C-D, −60%%±13%, p<0.05), whereas the catalytically inactive CDA mutant was ineffective. Moreover, that targeting CDA increases FANCD2 genomic instability markers in PDAC cells (Fig. 4C-D, +111%±7%, p<0.05). The latter was further validated in BxPC-3 (Extended Fig. 4A) and Capan-1 (Extended Fig. 4B) cells. Interestingly, markers of genomic stress such as remnants of incomplete replication can be inherited by daughter G1 cells and are sequestered to specific nuclear bodies that are colocalized with 53BP1. We found that CDA targeting in MIA PaCa-2 cells using shRNA significantly increases the number of G1 cells with 53BP1 bodies (Fig. 4E-F, +236%±14%, p<0.001). Collectively, these results strongly suggest that CDA participates in limiting genomic instability in PDAC cells.

Intrigued by these results, we extended our investigations to PDAC tumors. As shown in Fig. 5A, tumors with high levels of CDA expression demonstrate significant enrichment in activation of the ATR in response to replicative stress (Fig. 5A, normalized enrichment score = 1.72, p=0.01) and in the Cinsarc^27^ signatures of chromosome instability (Fig. 5B, normalized enrichment score =2.83 p<0.01). We next explored the aneuploidy score and the number of non-silent mutations per megabases of genomic DNA in PDAC tumors from TCGA that were classified as low or high for the expression of genes involved in genomic stability. As expected, POLQ, which is a low fidelity DNA polymerase that introduces various kinds of mutations during DNA repair, is enriched in PDAC tumors with high genomic instability (Fig. 5C-D, p<0.01). On the other hand, BRCA2 and DHODH expression levels were found to be unrelated with such markers of genomic instability of PDAC tumors (Fig. 5C-D). Remarkably, PDAC tumors with high levels of CDA demonstrate significantly higher aneuploidy scores (Fig. 5C, +69%±10%, p<0.01) and non-silent mutations per megabase (Fig. 5D, +43%±7%, p<0.005) as compared to tumors with lower CDA expression. Thus, we show that CDA levels are associated with genetically unstable PDAC tumours and could control genomic instability.

**Figure 5.**
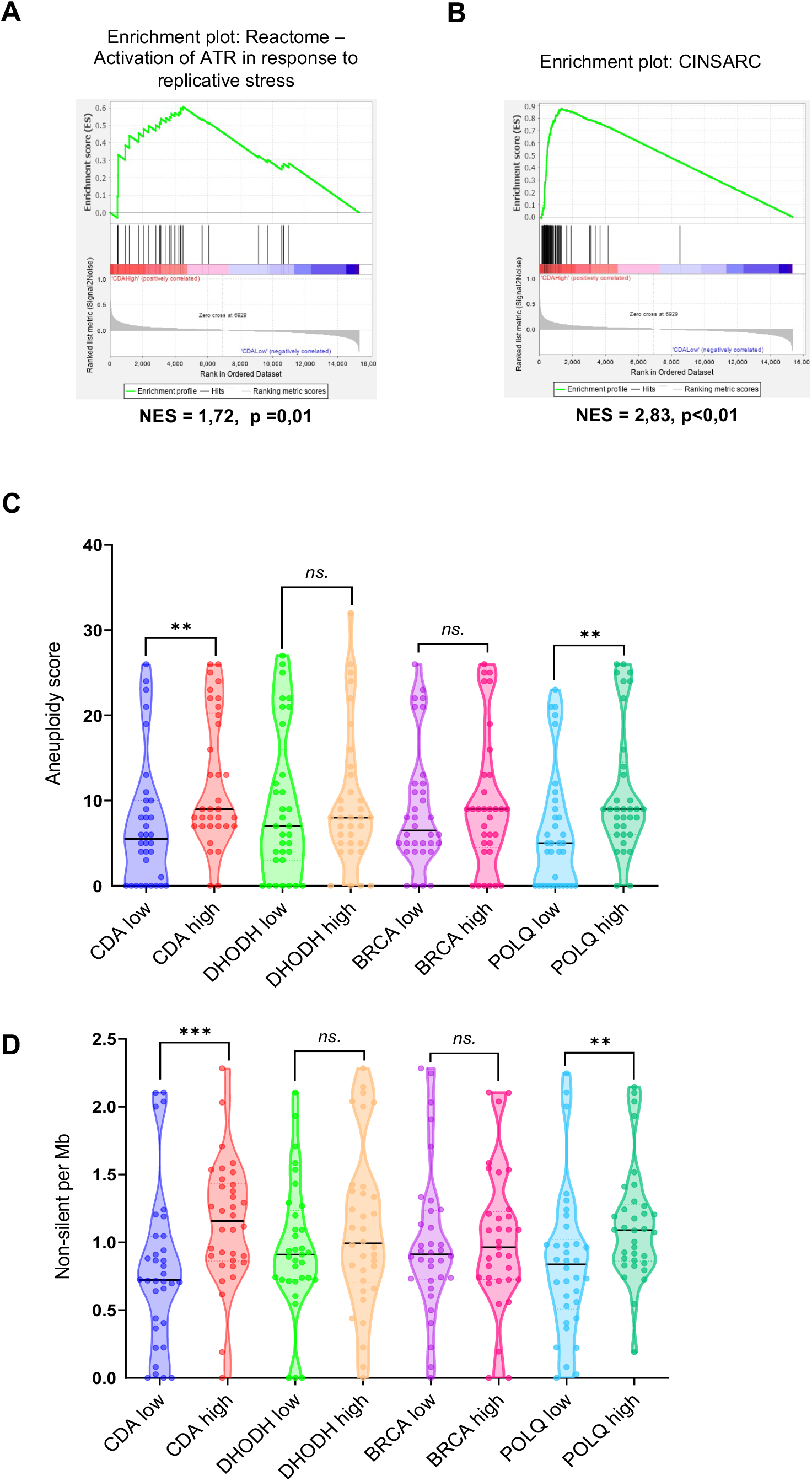
CDA expression is associated with genetic instability in PDAC tumors. Enrichment plots for **A**) Activation of ATR in response to replicative stress (Reactome – GSEA) and **B**) CINSARC signatures in TCGA PDAC samples expressing high or low CDA mRNA levels. **C.** Quantification of Aneuploidy score in TCGA PDAC samples with high and low levels of CDA, DHODH, BRCA and POLQ expression. Aneuploidy score is the total number of chromosome arms containing at least one variation of copy number in an arm, per sample (Taylor et al. 2018). **: p<0.01 (Wilcoxon-Mann-Whitney test). **D.** Quantification of the number of non-silent mutations per Mb in TCGA PDAC samples with high and low levels of CDA, DHODH, BRCA and POLQ expression. ***: p<0,001, **: p<0,01 (Wilcoxon-Mann-Whitney test) (TCGA PDAC samples: n=39 high CDA, n=38 low CDA, n=34 DHODH high, n=33 DHODH low, n=33 BRCA high, n=34 BRCA low, n=34 POLQ high and n=34 POLQ low).

### CDA drives resistance to DNA-damaging drugs in PDAC cells

Replication stress is starting to be rationally leveraged in cancer therapies, including for PDAC^8^, especially to better position traditional anticancer drugs, such as oxaliplatin, that act by increasing the replication stress within the tumor cell to induce cell death. Considering the new role of CDA we describe herein, we aimed to investigate whether CDA levels can predict DNA-damaging drug efficacy. We first explored the cancer cell line encyclopedia (CCLE) database and found that CDA expression significantly correlates with resistance to oxaliplatin (Fig. 6A, p=0.014) and to Topoisomerase I inhibitor irinotecan (Extended Fig. 6A, p=0.039) in a panel of 26 PDAC cell lines. We selected 4 cell lines with low (DAN-G, MIA PaCa-2, CAPAN-1, KP-4) or high (CAPAN-2, AsPC-1, BxPC-3, SU8686) CDA expression. As shown in Figure 6B, PDAC cells with high levels of CDA are 67-fold±0.3 (p<0.01) more resistant to oxaliplatin as compared to cells expressing low levels of the enzyme. We extended this observation to SN38, the active metabolite of CPT-11, a derivative of campthotecin^28^ that inhibits DNA topoisomerase I, Cisplatin, that forms DNA adducts, temozolomide, that deposits methyl groups on DNA guanine bases, and mitoxantrone, that causes single- and double-stranded disruptions after intercalating with the DNA molecule. PDAC cells with high CDA expression are 19-fold±0.2 (p<0.05), 9-fold±0.3 (p<0.05), 5.9-fold±0.3 (p<0.05) and 39-fold±0.15 (p<0.05) more resistant to SN38, cisplatin, temozolomide and mitoxantrone, as compared to cells expressing low levels of the enzyme, respectively (Fig. 6B). To obtain direct evidence of the role of CDA in the resistance to DNA-damaging drugs in PDAC cells, we treated MIA PaCa-2 cells expressing CDA or the catalytically inactive mutant of CDA (CDAE67Q) with 5μM camptothecin. Cells expressing luciferase were used as a control. As shown in Figure 6C, camptothecin shows similar antiproliferative activity in control cells and cells expressing a catalytically inactive mutant of CDA (Fig. 6C, −57%±5%, p<0.005). On the other hand, CDA expression significantly antagonizes the antiproliferative activity of the drug (Fig. 6C, −29%±2.5%, p<0.01), corresponding to a two-fold decrease in camptothecin efficacy.

**Figure 6.**
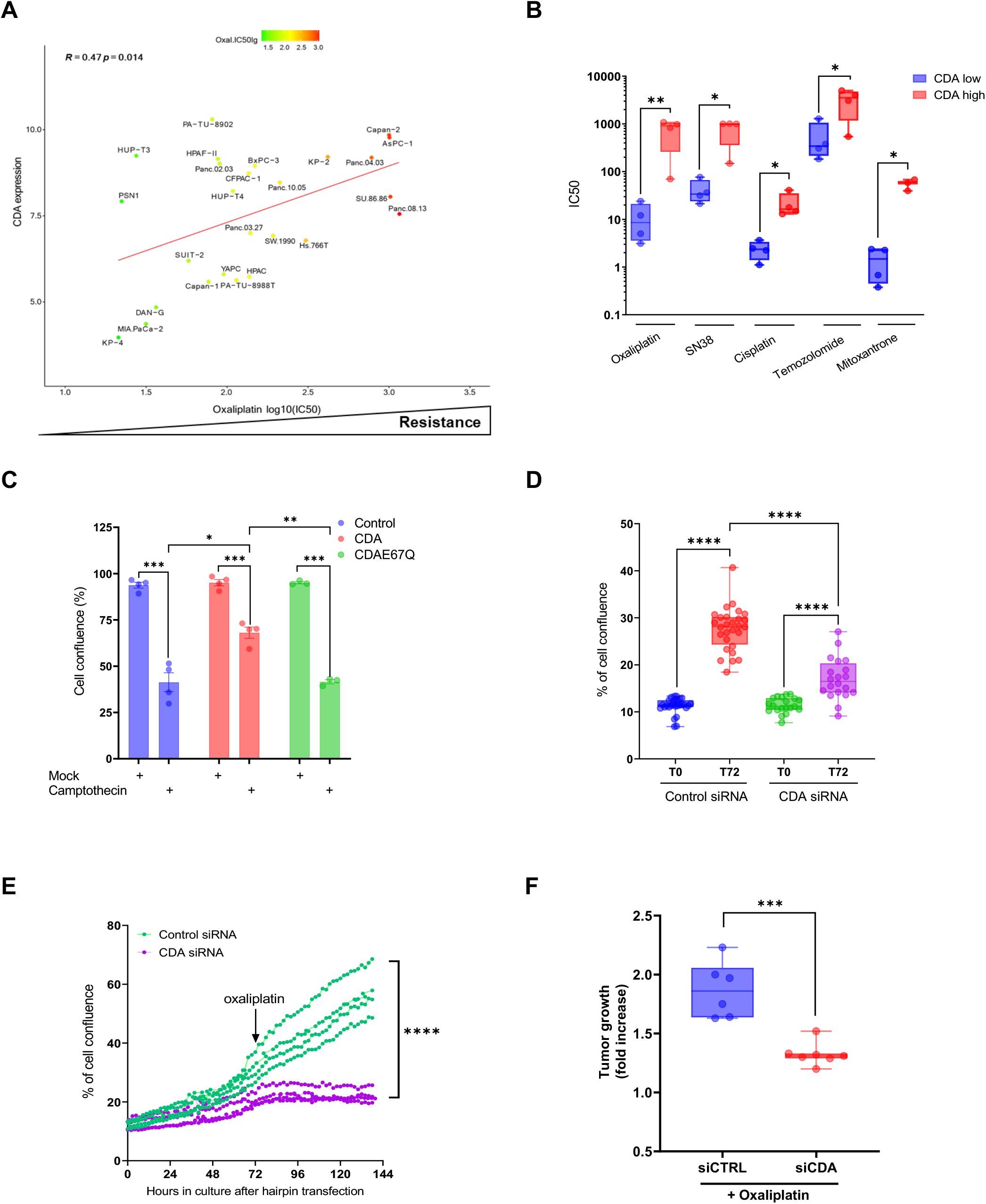
CDA drives resistance to replication stress-inducing drugs in PDAC cells. **A**. Correlation plot of CDA expression and log_10_IC_50_ for irinotecan in PDAC cell lines from CCLE. **B**. IC_50_ of irinotecan, SN38, cisplatin, temozolomide and mitoxantrone in PDAC cell lines with high and low CDA expression (n=4 for each). *: p<0.05, **: p<0.01 (unpaired t-test). **C**. Cell confluence (%) of MIA PaCa-2 control cells or overexpressing CDA and CDAE67Q catalytically inactive mutant and treated for 72h by 10μM campthotecin. *: p<0.05, **: p<0.01, ***: p<0.005 (unpaired t-test). **D**. Cell confluence (%) of PDAC051T cells depleted for CDA using siRNA 72 hours after transfection. **: p<0.01, ****: p<0.001 (unpaired t-test). **E**. Long term confluence follow-up of PDAC051T cells transfected with control or CDA siRNA, then treated by 10μM oxaliplatin ****: p<0.001 (unpaired t-test). **F**. Tumor growth (fold increase) of PDAC051T cells treated by siRNA targeting CDA and receiving oxaliplatin (n=7). Control tumors received control siRNA qnd oxapliplatin (n=6) ***: p<0.005 (unpaired t-test).

We next investigated whether CDA silencing could sensitize PDAC cells to DNA-damaging drugs. Thus, we used PDAC051T primary cells^29^, derived from a PDAC patient with known CDA expression, that were treated with siRNA targeting CDA for 72-hours. Control cells were transfected with a pool of control siRNA. We found that CDA silencing inhibits PDAC051T proliferation by 2-fold (Fig. 6D, p<0.0001), thus comfirming our finding that CDA is important for PDAC cell proliferation. We extended this finding to PDAC015T, deribed from another PDAC patient sample that also expressed high levels of CDA (Extended Fig. 6B, p<0.001). Next, PDAC051T cells were then treated by 10μM of the clinically-relevant drug oxaliplatin. As shown in Fig.6E and extended Fig. 6C, a combination of CDA silencing followed by oxaliplatin treatment significantly sentized PDAC ells to therapy and inhibited cell proliferation, as compared to primary cells transfected with control siRNA (−61%±5%, p<0.0001).

Further, we asked whether targeting CDA may represent a novel inerventional strategy to improve drug response in PDAC preclinical models, *in vivo*. Thus, PDAC051T were implanted in nude mice and transfected with siRNA targeting CDA. Mice with PDAC051T and transfected with control siRNA were used as control. Forty-eight hours later, mice were treated with oxaliplatin as previously described^29^. The combination of siRNA tranfection and oxaliplatin treatment was repeated three days later. Ten days later, mice were killed and tumors were measured and sampled. We found that tumors receiving siRNA targeting CDA and treated by oxliplatin showed a robust reduction in tumor growth as compared to control tumors (−29%±2%, p<0.005, Fig.6F and extended Fig.6D). Taken together, our results demonstrate for the first time that CDA expression is predictive of the therapeutic efficacy of DNA-damaging drugs in PDAC, and that CDA targeting sensitizes PDAC tumors to drugs inducing replication stress.

## DISCUSSION

Exploiting DNA replication stress has recently emerged as a promising strategy for cancer therapy^30^ and is beginning to be explored as an attractive target for PDAC management, a disease with no cure^7,9,10^. Still, the identification of the molecular mechanisms involved in the dependency of tumor cells on the replication stress response is key to devise future and selective therapies centered on enhancing both endogenous and drug-induced replication stress.

In this work, we identify that CDA, a key protein from the pyrimidine salvage pathway, is overexpressed in the most lethal PDAC tumors, that CDA silencing provokes tumor cell death by apoptosis, and it strongly inhibits experimental tumor growth. To our knowledge, this is the first description that CDA is essential to PDAC tumor growth, independently of its role in tumor resistance to pyrimidine-based therapies^11^. Others have identified that PDAC cells are dependent on *de novo* pyrimidine production driven by dihydroorotate dehydrogenase (DHODH)^31,32^. However, targeting DHODH would not be selective of PDAC cells, as we found that DHODH is equally expressed at low levels in normal and tumoral pancreatic tissues (personal data and TCGA analysis).

*In silico* exploration of PDAC patient samples indicates that CDA expression is strongly associated with DNA replication signature. In PDAC cell lines, we show that CDA promotes DNA replication and reduces endogenous replication stress. Interestingly, these effects are independent of cell proliferation (data not shown). Importantly, we report here that the reduction of CDA levels in PDAC cell lines slows fork progression and increases γ-H2AX levels in cells that are checkpoint and DNA damage repair proficient. Considering that replication nucleotide shortage is one of the main promoters of replication stress and that catalytically dead CDA mutants fail to reduce the level of DNA breaks in PDAC cells, our results strongly suggests that CDA controls PDAC cells replication stress by recycling pyrimidines from nucleotide degradation through the salvage pathway. This occurs upstream of the reduction of ribonucleoside diphosphates to deoxyribonucleoside diphosphates by the ribonucleotide reductase (RNR). Thus, our results may help revisit the central position of RNR in controlling DNA replication in PDAC cells. Notably, we show in this study that CDA, generally considered a cytoplasmic enzyme^11^, is nuclear and locates at the replication fork, but the mechanism involved is currently unclear. In proteomic databases, CDA is documented to co-eluate with Minichromosome Maintenance Complex Component 2 (MCM2)^11^, but we could not see this association in PDAC cells (data not shown). Hence, our findings may revive the “dNTP channelling hypothesis” based on the existence of a functional complex, known as ‘replitase’^33^, that concentrates dNTP in the environment of replication forks for maximal efficacy. Replitase is thought to contain enzymes that are involved in nucleotide precursors synthesis and DNA replication, such as RNR^34^, NDP-kinase (NDPK) and DNA polymerase α (Pol α)^35^.

Interestingly, as deoxyribonucleotides are detected in newly synthesized DNA much faster than in the dNTP pool^36^, this strongly suggests that the salvage pathway may also actively participate to channel dNTPs to the fork environment. Further studies are underway to characterize the interactome of CDA in these cells, and to determine whether CDA belongs to the replitase complex.

Although replication stress contributes to cancer development by promoting genomic instability, it slows down cell proliferation and activates anticancer barriers leading to apoptosis or senescence^37^. To proliferate, cancer cells must therefore bypass these barriers, while avoiding severe replicative defects that are incompatible with cell survival. It is generally believed that cells adapt to replication stress by modulating the intensity of the ATR-CHK1 checkpoint response. Here, we found that CDA dampens endogenous replication stress in PDAC cells and is overexpressed in genetically unstable tumors. In addition, we found that CDA expression is specifically elevated in primary cells from the squamous molecular subtype, which is strongly associated with replication stress^8^. Therefore, it is likely that the adaptation to replication stress mediated by increased CDA expression occurs at the expense of genome integrity and would therefore promote cancer progression. Another important question raised by our study concerns the mechanism by which cancer cells overexpress CDA. Oncogene activation drives replication stress, particularly through RAS and MYC signalling, both of which are prevalent molecular features of PDAC^38^. While this field is currently unexplored in PDAC, considering the new role of CDA we describe here, it is tempting to speculate that CDA is expressed as a result of oncogene-induced replication stress to sustain PDAC tumors’ growth.

Our work may have important implications for the treatment of patients, as it could influence future studies. With a handful of mechanism-based therapies, the future of PDAC treatment lies in rational therapy combinations for durable tumor control in patients. Our study provides a strong scientific rationale for using CDA expression as a marker of resistance to DNA-damaging drugs used in routine clinical practice such as SN38 and oxaliplatin. In addition, we show that targeting CDA strongly increases tumor replicative stress and sensitizes to oxaliplatin treatment, both *in vitro* and *in vivo*. Interestingly, Dreyer et al have recently proposed an elegant matrix for therapeutic decision using DDR agents and/or cell cycle checkpoint inhibitors based on the DDR proficiency and the level of replication stress of PDAC cells^8^. Unfortunately, DDR-proficient PDAC cells with low replication stress that account for half of samples, respond to neither class of agent. Thus, interventional studies aimed at targeting CDA have the potential to raise the level of replication stress in PDAC tumors in combination with ATR or WEE1 inhibitors, which can be further combined with PARP inhibitors in the context of DDR deficiency.

## MATERIALS AND METHODS

### Cell lines and culture conditions

MIA PaCa-2, BxPC-3, Capan-1, Hela S3 human cancer cells were obtained from ATCC and cultured in Dulbecco Modified Eagle Medium (DMEM) 4.5g/L glucose (for MIA PaCa-2) or Roswell Park Memorial Institute medium (RPMI) (BxPC-3 and Capan-1) containing 10% Fetal Bovine Serum (FBS), 100 IU/mL penicillin, 100μg/mL streptomycin, and 250ng/mL fungizone (AB). Cells were incubated at 37°C with 5% CO_2_. Pancreatic cancer patient-derived cell lines (PDCL) PDAC015T and PDAC051T were cultivated as described before^29^. Cells were incubated at 37°C with 5% CO_2_.

### Mouse models

Experimental procedures performed on mice were approved by the ethical committee of INSERM CREFRE US006 animal facility and authorized by the French Ministry of Research: APAFIS#3600-2015121608386111v3. MIA PaCa-2 (2×10e6) stably expressing luciferase and shRNA targeting CDA following lentiviral delivery were engrafted into the pancreas of SCID CB17 mice as previously described^39^. MIA PaCa-2 cells expressing random hairpins were used as control. Tumor progression was monitored once per week by luminescence after intra peritoneal injection of Rediject D-Luciferin (Perkin Elmer). Measurement of luminescence was assessed with IVIS Spectrum (Perkin Elmer). For interventional studies, 1×10e6 PDAC015T cells resuspended in matrigel were injected subcutaneously in atymic mice. Tumors were measured with a caliper and tumor volume was calculated using the formula V = (Width^2^ × Lenght)/2. Two weeks later, tumors ranging from 150 to 350 mm^3^ were randomized in two groups (day 0). Tumors were injected intratumorally at days 0 and 5 with 4μg of siRNA pool targeting CDA (Dharmacon) using *in vivo* JetPEI (N/P=6) according to the manufacturer’s recommendation (Polyplus). Tumors receiving control siRNA pool (Dharmacon) were used as control. At days 2 and 7, mice received 5mg/kg oxaliplatin I.V. Mice were sacrificed at day 10.

### Patient cohorts and clinical sample processing

We used two patient cohorts for CDA expression analysis in PDAC and normal adjacent tissues. The first one includes pancreatic tissue samples obtained from patients receiving pancreatic surgery following the policies and the practices of the facility’s ethical committee at the Centre Hospitalo-Universitaires (CHU) of Toulouse and Bordeaux, and the Cancéropole Grand Sud-Ouest (France), as stated before^40^. All patients gave their informed written consent. Histopathology faculty selected cancerous pancreatic tissue with matched adjacent tissue. RNA was extracted and processed as stated before^40^. The second cohort was established from pancreatic carcinoma tissues and normal adjacent pancreatic parenchyma collected from 48 patients who had undergone surgical resection for ductal adenocarcinoma of the pancreas between 2003 and 2011 at the Bellvitge Hospital, Barcelona, Spain^41^. The study was approved by the Ethical Committee of University Hospital of Bellvitge CEIC 02/04 and written informed consent was obtained from all patients for the use of their tissues.

### Plasmid cloning

For silencing studies, lentiviral vectors encoding for shRNAs sequences targeting CDA (CCGGCATGAGAGAGTTTGGCACCAACTCGAGTTGGTGCCAAACTCTCTCATGTTTTTG) were cloned into pLKO.1 puromycin vector and obtained from Sigma. Vectors encoding for random shRNA (Sigma) were used as a control. For overexpression studies, we generated an open reading frame (ORF) for CDA by PCR that was subcloned in pDonor221 (Invitrogen). CDA E67Q catalytically inactive mutant^42^ was generated by site directed mutagenesis according to the manufacturer instructions (New England Biolabs). Lentiviral expression constructs were obtained by cloning CDA and CDAE67Q ORF into pCMV blasticidin DEST 706-1 vectors (Addgene) using the Gateway strategy (Invitrogen). Luciferase was used as a control. All constructs were sequence verified. Lentiviral particles were produced in 293FT cells as previously described^35^ and quantified for p24 presence using INNOTEST HIV Antigen mAb (Fujirebio). For lentiviral transduction of shRNAs and ORF-expressing constructs, 250ng p24/50.000cells were used in Opti-MEM medium (ThermoFisher Scientific) containing 4μg/mL protamin sulfate, as previously described^35^. Transductions were performed overnight, the medium was changed the next day, and transduced cells were selected with 5μg/mL 2 days later. Cell cultures were maintained as pools and certified mycoplasma-free (Lonza).

### Proliferation assay

For proliferation studies, 6×10e3 cells were seeded in 96-well plates in 100μl of complete medium. Medium was changed and cells were treated 24h after seeding. Confluence was quanitfied non-invasively using the Incucyte Zoom (Sartorius). For silencing studies, siRNA smartpools were purchased from Dharmacon. Cells were transfected with 20nM siRNA targeting CDA using JetPrime (Polyplus) as per the manufacturer recommendation. Control cells were transfected with control siRNA (Dharmacon). Three days later, cells were treated or not with 10uM oxaliplatin and cell confluence was monitored as previously described.

Colony formation assays were performed in 6-well plates, with 250 cells initially seeded. Colony presence was revealed at day 10 by cold methanol staining and Crystal violet coloration. Density was measured with Image lab software (BioRAD).

### RNA isolation and gene expression analysis

Total RNA was extracted with RNeasy kit (Qiagen) according to the manufacturer’s instructions; quality and quantity were measured on a Nanodrop (Thermo Scientific). cDNA synthesis was performed with Revertaid H minus kit (Thermo Scientific). cDNA expression analysis was performed using Sybr Green (Bio-RAD) quantitative real time PCR on a StepOne (ThermoFisher Scientific). CDA forward: GGGG ACA AGT TTG TAC AAA AAA GCA GGC TAT GGC TAT GGC CCA GAA GCG T. CDA reverse: GGGG AC CAC TTT GTA CAA GAA AGC TGG GTT CAC TGA GTC TTC TG.

### Western Blot

Total cell lysate protein extraction for immunoblot analysis was performed using RIPA buffer (Biotech) (Tris-HCl 50mM pH=8, NaCl 150mM, NP40 0,5%, sodium deoxycholate 0,5%, SDS 0,1%) in the presence of protease inhibitors (Sigma). Extracts were separated by SDS-PAGE under reducing conditions, transferred to a nitrocellulose membrane and analysed by immunoblotting for cleaved PARP (bethyl 9541), Cleaved caspase-3 (Cell signalling #9664, 1:1000), P-Chk1 S345 (Cell signaling #2348, 1:1000), Chk1 (Cell signaling #2360, 1:1000), CDA (abclonal A13959, 1:1000), FLAG, (Sigma Aldrich F1804, 1:5000), GAPDH (Cell signaling #5174, 1:5000), actin β (santa cruz sc-47778, 1:2000), HSP90 (Cell signaling #4874, 1:2000), SP1 (Santa cruz sc-59 1:5000), 4EBP1 (Cell signalling #9644 1:5000), PCNA (Sigma Aldrich P8825), MCM7 (Abcam ab2360). Appropriate HRP conjugate secondary antibodies were from Promega. Signal was detected using the ECL system (BioRAD) according to the manufacturer’s instructions. Densitometry of the blots was done using Chemidoc (BioRAD) and image Lab software.

Cytoplasmic/nuclear extract isolation was performed as following: cell pellet was resuspended in 10mM TRIS, pH 7.4 containing 1.5mM MgCl2, 5mM KCl, 0.5mM dithiothreitol, 0.5%NP40 and 0.5mM phenylmethylsulfonyl fluoride complemented with protease inhibitors and incubated on ice for 10min. The mixture was centrifuged for 15min at 4°C at 2000 rpm. Supernatant was collected (cyplasmic fraction) and the pellet was washed and centrifuged twice with the same buffer as previously. The pellet was then resuspended in 20mM TRIS, 0.025% glycerol, 1.5mM MgCl_2_, 0.5mM phenylmethylsulfonyl fluoride, 0.2mM EDTA, 0.5mM dithiothreitol and 0.4M NaCl complemented with protease inhibitors and incubated on ice for 15min. The mixture was centrifuged 20min at 4°C at 12000rpm and supernatant was collected (nuclear fraction).

### Indirect immunofluorescence protocol

Cells were seeded in 6-well or 12-well plates, containing coverslips, in 1-3mL of complete medium. Medium was changed and cells were treated 24h after seeding. For the study of cells in S-phase, EdU 10μM (5-ethynyl-2’-deoxyuridine, ThermoFisher Scientific) diluted in fresh medium was added in the well for 10 minutes (MIA PaCa-2), 15 minutes (Capan-1) or 20 minutes (BxPC-3). Then, cells were preextracted with 0.2% triton 3 minutes, fixed in 4% PFA 15 minutes and permeabilized with 0.5% Triton X-100 for 20 minutes. For revealing EdU, the Click-iT EdU Imaging kits protocol (ThermoFisher Scientific) was followed. Samples were blocked with PBS 5% BSA. Primary antibodies were mouse anti-phospho-Histone (Ser139) (JBW301, Milipore 05-636, 1:1000), rabbit anti-phospho-53BP1 (S1778) (Cell Signaling #2675, 1:1000), rabbit anti-phospho-RPA2 (S4-S8) (Bethyl A300-245A, 1:500), rabbit anti-RPA70 (Cell Signaling #2267, 1:500), rabbit anti-FANCD2 (NOVUSBIO NB100-182, 1:1000) and rabbit anti-53BP1 (Bethyl A300-272A, 1:2500). DNA was stained with 4,6-diamidino-2-phenylindole (DAPI) and images acquired as followed. Images acquires on Nikon DS-Qi2 fluorescence microscope and ImageJ and CellProfiler were used to quantify the number of foci and staining intensity per nucleus. For each condition, at least 500 cells were measured. For quantification in early mitotic cells, at least 50 prophase and pro-metaphase cells were counted.

### DNA fibre assay

Cells were seeded in Ø6cm petri dishes in 3mL of complete medium. Cells were incubated with 50μM IdU and 100μM CldU for 30 minutes at 37°C, with washing between the two pulses. Cells were harvested, resuspended (0.5 10^6^ – cells/mL in PBS) and 2μL were spotted onto microscope slides after what were added 7μL of lysis buffer (200mM TriHCl pH7.4, 50mM EDTA 0.5M, 0.5% SDS). Glass slides were tilted and dried overnight, DNA spreads were then fixed in ethanol/acetic acid (3:1) 20 minutes at −20°C. Samples were incubated in Pepsin Buffer (0.5mg/mL Pepsin, 30mM HCl) at 37°C for 20 minutes, denaturated in HCl 2.5M at 37°C for 45 minutes and blocked in PBS 1% BSA 0.1% tween-20. DNA fibers were incubated with mouse-FITC-anti-bromodeoxyuridine (detects IdU, B44, Becton Dickinson, 1:50) and rat-anti-bromodeoxyuridine (detects CldU, abcam ab6326, 1:100), at 37°C for 1 hour and then with anti-mouse IgG AlexaFluor 488 (Invitrogen A11029, 1:200) and anti-rat IgG AlexaFluor 555 (Invitrogen A21094, 1:200), at 37°C for 1 hour. Images were acquired on Nikon DS-Qi2 fluorescence microscope and Axio Observer Z1 Zeiss and ImageJ was used to quantify the length of DNA tracks in μm. In each condition, at least 100 fibres were measured.

### iPOND assay

EdU-labeled sample preparation. MIA PaCa-2 cells overexpressing CDA-FLAG (^~^2.10^8^ cells per condition) were incubated with 10μM of EdU for 15 minutes. For pulse-chase experiments with thymidine, EdU-labeled cells were washed once with warm media to remove the EdU and then incubated with 10μM thymidine for 2 hours. Next, cells were cross-linked in 2% PFA for 15 minutes, quenched using 0.125M glycine and washed with three times with PBS. Collected cell pellets were frozen at −80°C. Cells were permeabilized with 0.5% triton for 30 min and click chemistry was used to conjugate biotin-TEG-azide (Eurogentec) to EdU-labelled DNA in PBS containing 10 mM sodium Ascorbate, 10 μM biotin-TEG-azide, 2 mM CuSO_4_. Cells were re-suspended in lysis buffer (10 mM Hepes-NaOH; 100 mM NaCl; 2 mM EDTA PH8; 1 mM EGTA; 1 mM PMSF; 0.2% SDS; 0.1% Sarkozyl) and sonication was performed using a Qsonica sonicator with the following settings: 30% power, 20 sec constant pulse and 50 sec pause for a total sonication time of 5 min on ice with water. Lysates were centrifuged at 13,200 rpm for 10 mins at RT. Supernatants were normalized by DNA quantification using a nanodrop device. Biotin conjugated DNA-protein complexes were captured using overnight incubation with magnetic beads coated with streptavidin (Ademtech). Captured complexes were washed with lysis buffer and 500 mM NaCl. Proteins associated with nascent DNA were eluted under reducing conditions by boiling into SDS sample buffer for 30 min at 95°C and analyzed by Western blot.

### RNAseq data analysis

RNA-seq data was first converted from Illumina format to fastq using bcl2fastq (https://support.illumina.com/sequencing/sequencing_software/bcl2fastq-conversion-software.html). Quality control was performed with [fastqc](https://www.bioinformatics.babraham.ac.uk/projects/fastqc/) and adaptor sequences were trimmed using [Trimmomatic](http://www.usadellab.org/cms/?page=trimmomatic). Quality control with fastqc was run again after adaptor removal. Gene expression was quantified at the transcript level by [Salmon](https://salmon.readthedocs.io/en/latest/salmon.html). Import of Transcript-level abundances and counts from Salmon and name conversion to [HUGO](https://www.genenames.org/) gene symbols was performed with the [tximport](https://bioconductor.org/packages/release/bioc/html/tximport.html) R package. Differential expression analysis was performed with DeSeq2 using CTRL samples as the reference for the sign of the log2FoldChange.

### Functional enrichment

Enrichment analyses were performed using GSEA [Hallmarks](https://www-sciencedirect-com.proxy.insermbiblio.inist.fr/science/article/pii/S2405471215002185?showall%3Dtrue%26via%3Dihub) and Reactome and most other pathways were extracted from [MSigDB](https://doi-org.proxy.insermbiblio.inist.fr/10.1093/bioinformatics/btr260). No genes were removed during the analysis.

### TCGA

HTSeq counts for pancreas were downloaded from the gdc portal. Curated pancreas samples were filtered based on information from^18^. Samples were classified according to low (less than second quartile), normal (between second and third quartile) and high (higher than third quartile) values of CDA expression. Differential expression analysis comparing low CDA vs high CDA samples was performed with DeSeq using low samples as the reference for the sign of the log2FoldChange. A similar procedure was used to define the low/high DHODH/BRCA/POLQ sample sets.

### Instability signatures and metrics

The CINSARC signature gene list was obtained from^27^. Aneuploidy score and Mutational Load score (number of mutations per Mb) data was downloaded from the gdc portal.

### Statistical analysis

Unpaired Student’s *t*-or Wilcoxon-Mann-Whitney tests were used to determine the statistical significance of differences between two groups using GraphPad Prism 9 software with the default settings. Methods of statistical analysis are indicated in the figure captions. Values are presented as \**P* <0.05, \*\**P* <0.01 and \*\*\**P* < 0.005 and **** *P* <0.001. Error bars are s.e.m. unless otherwise stated. The experiments were performed with a sample size *n* greater than or equal to three replicates, and the results from representative experiments were confirmed in at least two independent experiment repeats and multiple cell lines. For the animal studies, female mice were used in an age-matched manner. Sample sizes (*n* ≥ 5 mice) were calculated using a *t*-test for two-group independent samples (ensuring that a power of 0.8 was reached) and a significance level of 0.05. *P* < 0.05 was considered statistically significant. When monitoring tumor growth, the investigators were blinded to the group allocation during bioluminescence imaging but were aware of group allocation when assessing the outcome. For the other experiments, sample size was determined empirically based on our preliminary or previous studies; experiments were not randomized; and the investigators were not blinded to allocation during the experiments or outcome assessment. No data were excluded from the analyses.

## Acknowledgements

The authors would like to thank the financial support of Fondation Toulouse Cancer Santé, Inserm, Région Occitanie, Université Paul Sabatier UT3 and Ligue Nationale Contre le Cancer. The authors would like to thank Ms Emilie Martin for technical assistance and Mr Jacobo Solorzano for data analysis. The authors would like to thank Pr Jerôme Cros and Dr Jérôme Torrisani for helpful discussions.

---

**Extended Figure 1.**
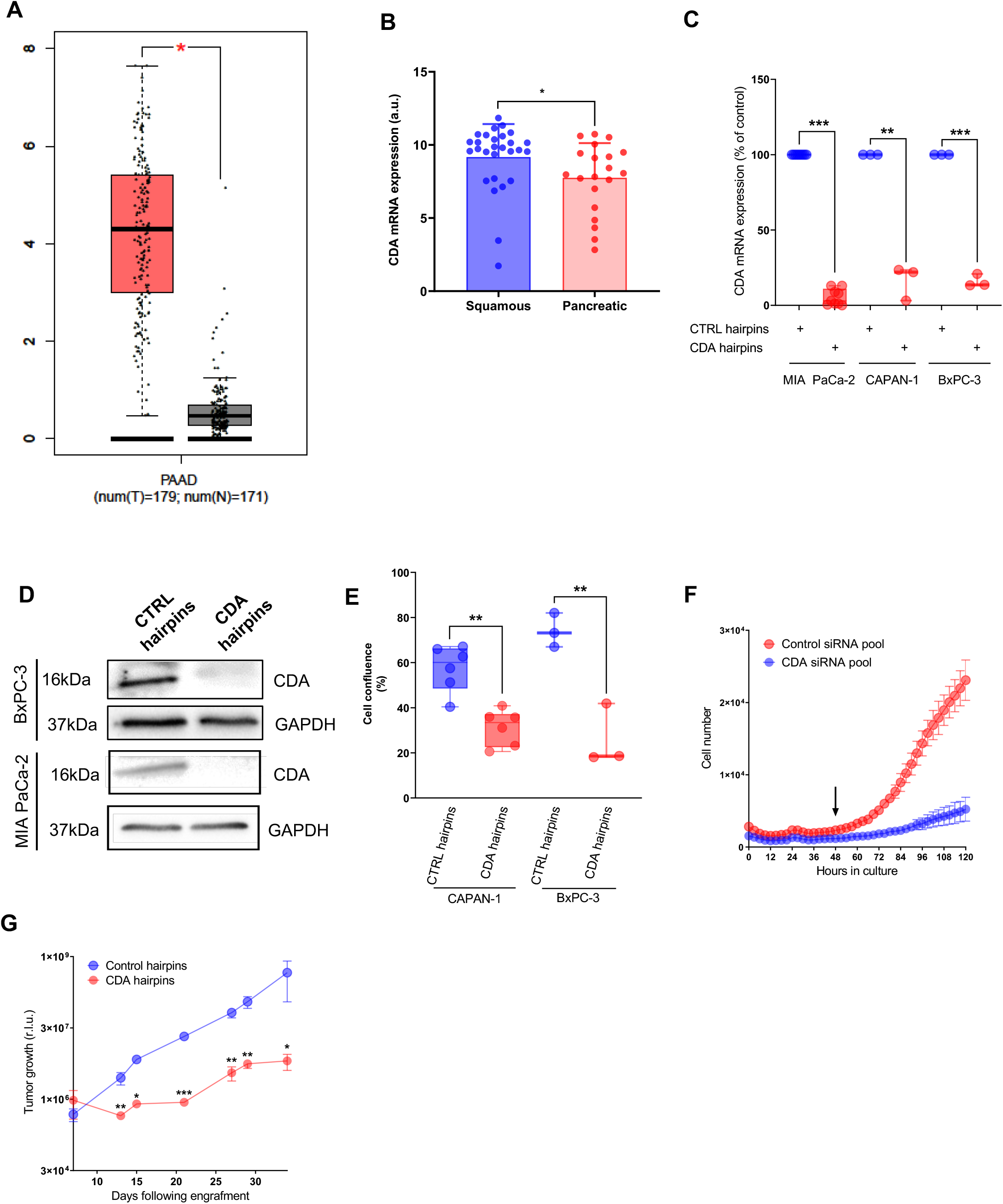
CDA is overexpressed in PDAC tumors and essential for cell proliferation and tumor growth. **A**. CDA expression (mRNA) in pancreatic tumors (red, n=179) and normal pancreas (grey, n=171) of TCGA PDAC samples. *: p<0,05 (unpaired t-test). **B**. CDA mRNA expression in primary cells from squamous and pancreatic molecular subtype. *: p<0.05. **C**. CDA mRNA expression in MIA PaCa-2 receiving CDA hairpins vs control cells (control hairpins). Mean of nine independent pools of transduction. ***: p<0.001 (unpaired t-test). **D**. Western Blotting of CDA in MIA PaCa-2 cells receiving CDA hairpins vs control cells (control hairpins). Representative of at least nine independent pools of transduction. Long term follow-up of cell confluence (%) of Capan-1 cells (**E**) transduced with CDA hairpins and control hairpins and (**F**) of BxPC-3 transfected with CTRL siRNA or CDA siRNA. **G**. Mean quantification of the tumor growth of MIA PaCa-2 cells expressing CDA hairpins as compared to control cells, and assessed by luminescence (n=6 per group). *: p<0.05, **: p<0.01, ***: p<0.001 (unpaired t-test).

**Extended Figure 2.**
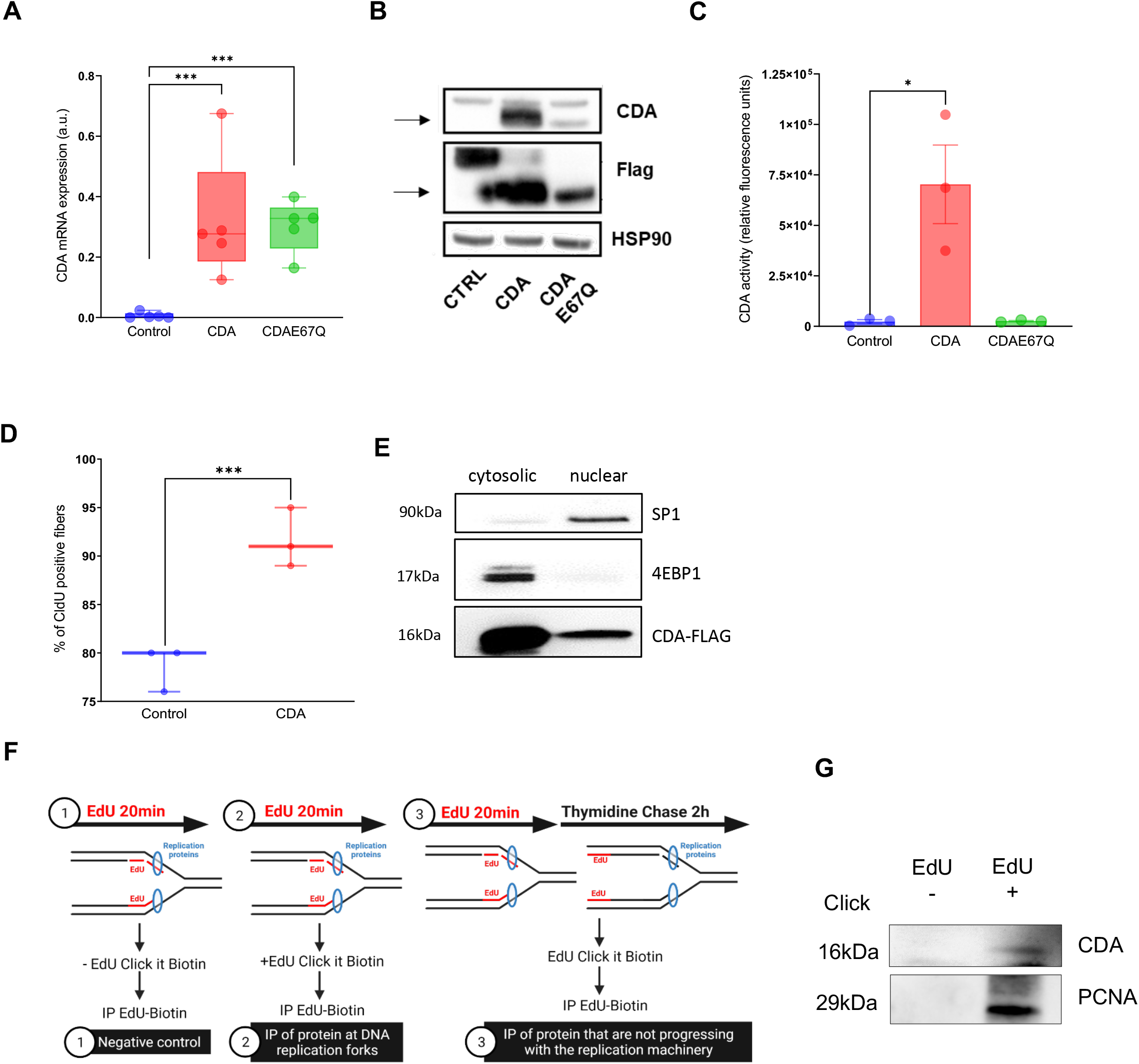
CDA increases replication fork speed and restart efficiency in PDAC cells. **A**. CDA expression (mRNA) in cells overexpressing control, CDA or CDAE67Q. Representative of five transduction pools. ***: p<0.001 (paired t-test). **B**. Western Blotting for CDA and FLAG in cells overexpressing control, CDA–FLAG or CDAE67Q-FLAG cells. HSP90 is used as a loading control. Representative of five transduction pools. **C**. CDA activity (relative fluorescence units) measured by CDA activity assay (fluorometric assay) in MIA PaCa-2 cells overexpressing luciferase (control), CDA or CDAE67Q. Representative of three transduction pools. *: p<0.05 (unpaired t-test). **D**. Percentage of CldU positive fibers following HU treatment, in cells overexpressing CDA versus cells overexpressing luciferase (control). Representative of three independent transduction pools. ***: p<0.001 (unpaired t-test). **E**. Western Blotting of CDA-FLAG, 4EBP1 and SP1 in cytosolic and nuclear fractions of MIA PaCa-2 cells overexpression FLAG-CDA cells. Representative of three independent experiments. **F**. Schematic procedure of iPOND experiment. **G**. IPOND experiment in Hela S3 cells. PCNA is used as control. Results are representative of 3 independent experiments.

**Extended Figure 3.**
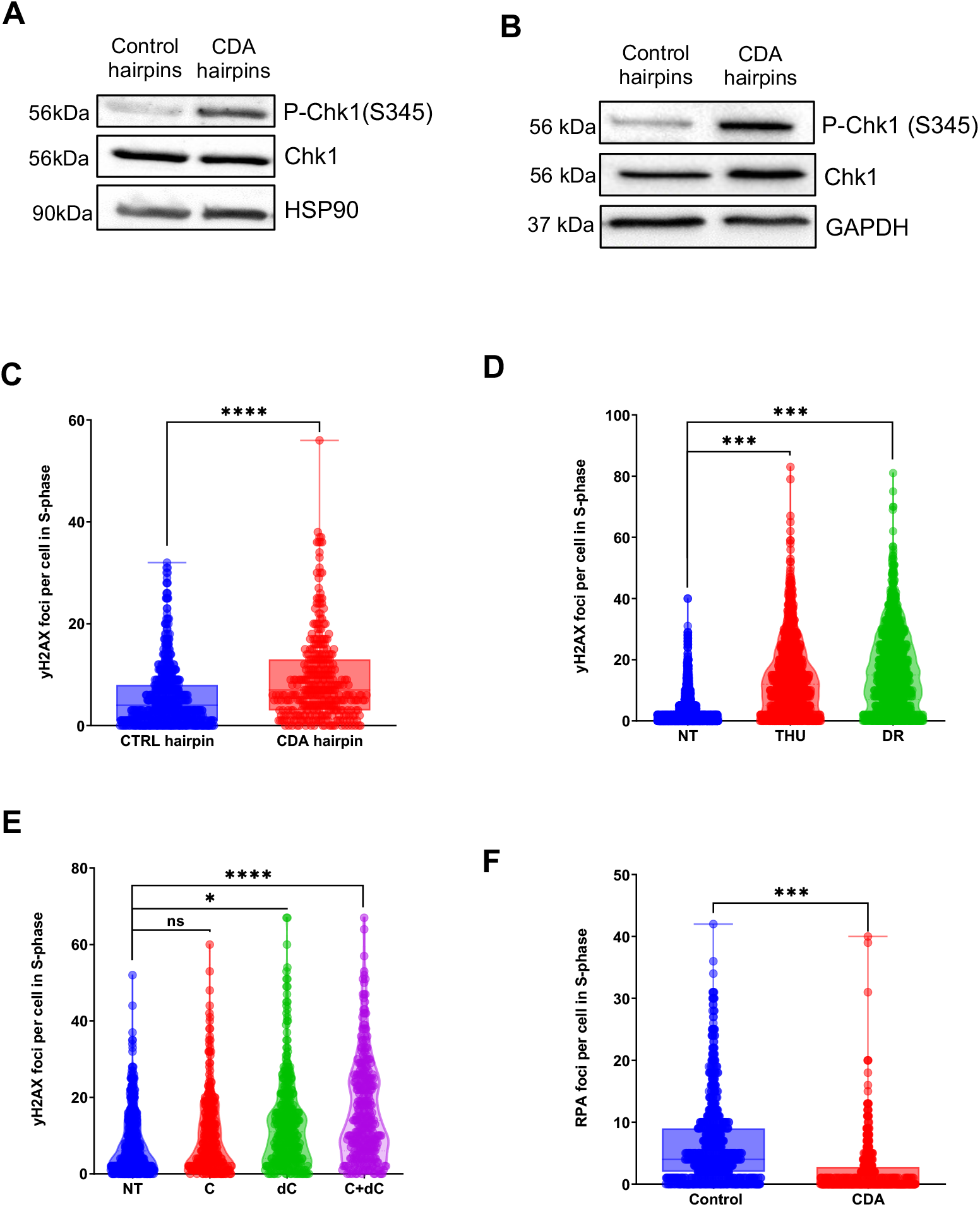
CDA controls replication stress levels of PDAC cells. Western Blotting for P-Chk1 (S345) and Chk1, HSP90 and GAPDH were used as loading control, in (**A**) control Capan-1 cellls and Capan-1 cells expressing CDA hairpins and in (**B**) control BxPC-3 cells and BxPC-3 cells expressing CDA hairpins **C.** Quantification of the number of y-H2AX foci per cell in S-phase in Capan-1 cells depleted of not for CDA (CDA hairpins, control hairpins). Results are representative of three independent transduction pools. ****: p<0,0001 (Wilcoxon-Mann-Whitney test). Quantification of the number of y-H2AX foci in S-phase cells in MIA PaCa-2 treated with (**D**) 100μM of CDA inhibitors THU and DR for 72 hours or (**E**) 1mM Cytidine (C), Deoxycytidine (dC) or both for 72 hours. Results are representative of two independent experiments. *: p<0.05, ***: p<0.001, ****: p<0.0001 (Wilcoxon-Mann-Whitney test). **F.** Quantification of RPA foci per S-phase cells (EdU+) (at least 500 cells) in MIA PaCa-2 cells overexpressing CDA versus control cells. Results are representative of three independent transduction pools. ***: p<0.001 (Wilcoxon-Mann-Whitney test).

**Extended Figure 4.**
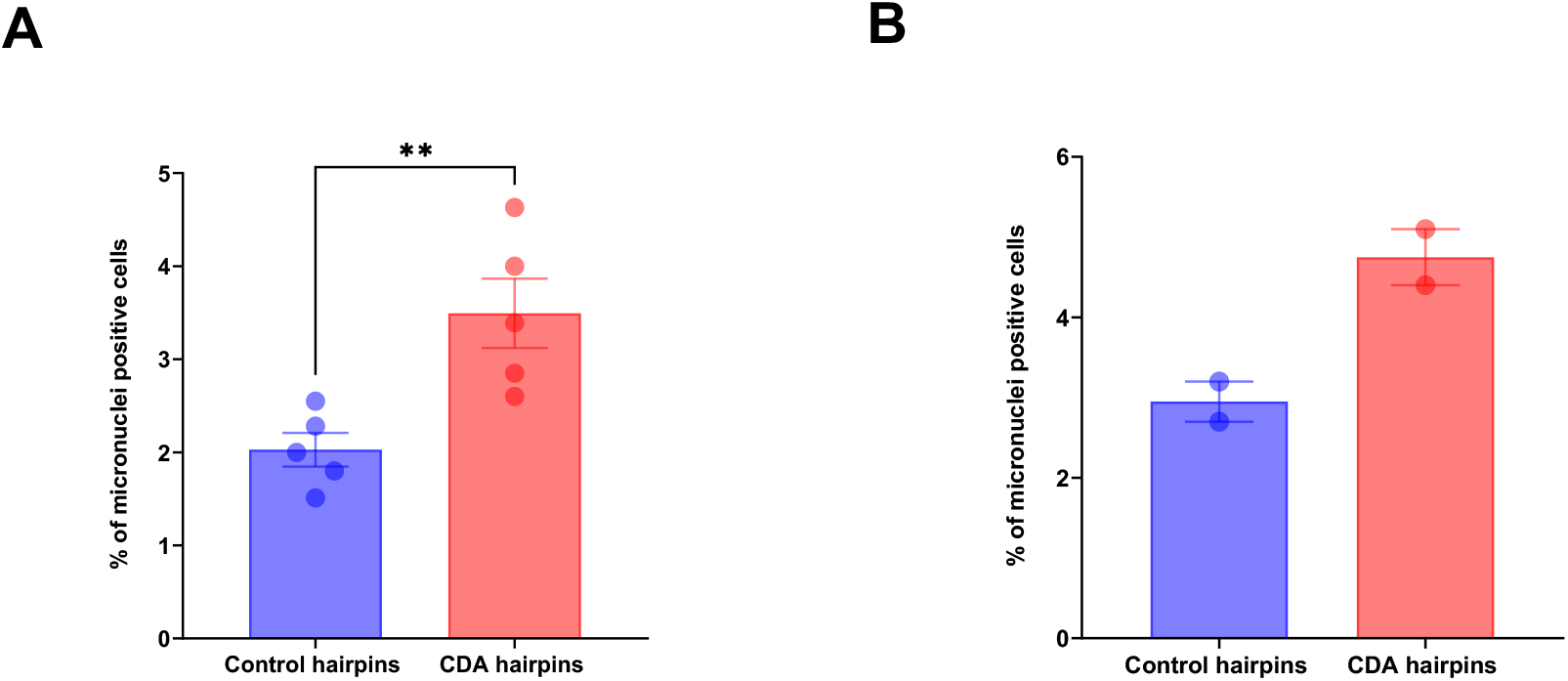
CDA controls genomic instability of PDAC cells. Quantification of the percentage of Capan-1 (**A**) or MIA PaCa-2 cells (**B**) depleted for CDA with at least one micronucleus as compared to control and. Representative of five independent experiments for Capan-1 and two independent experiments for MIA PaCa-2. **: p<0,01 (paired t-test).

**Extended Figure 6.**
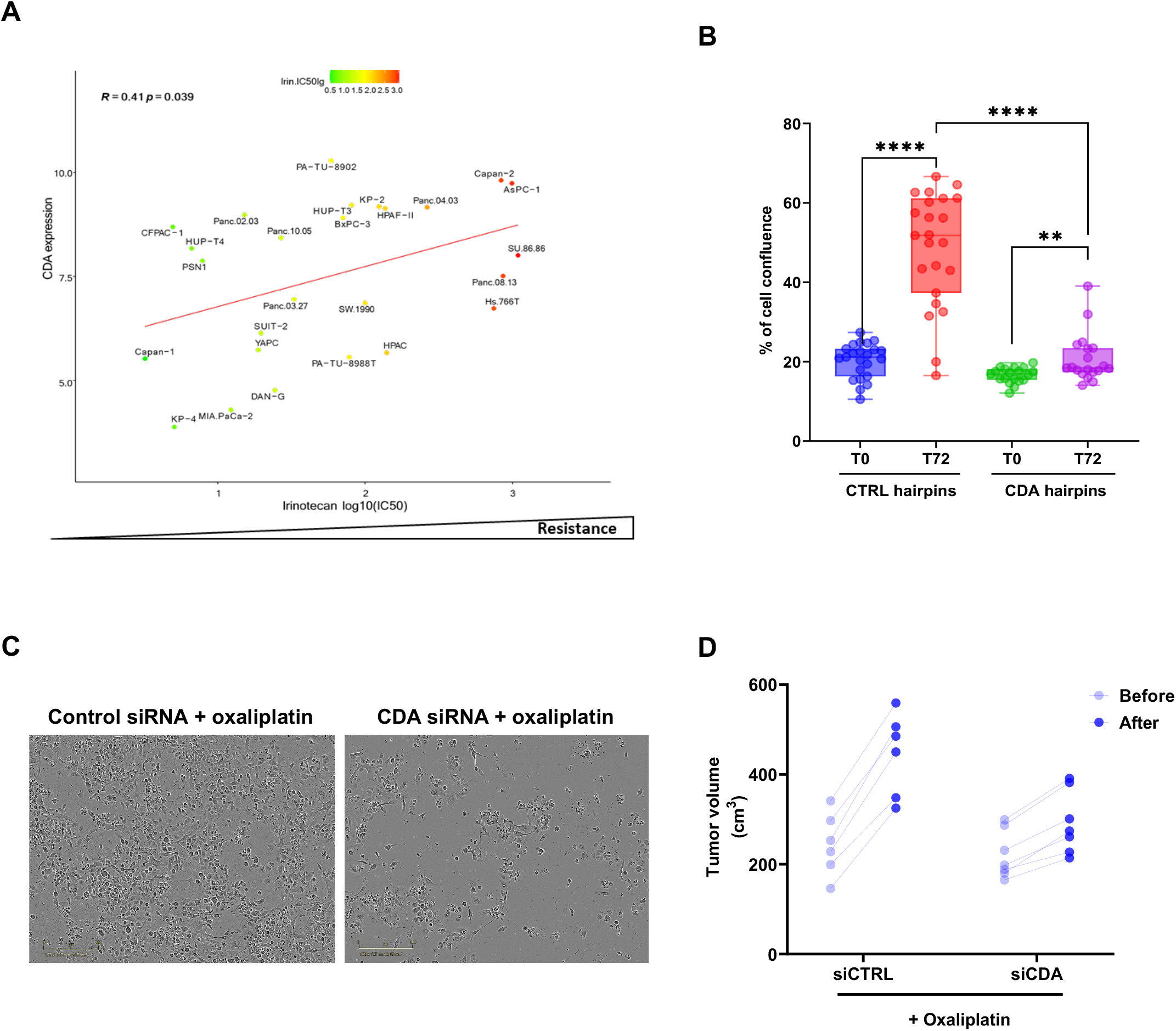
CDA drives resistance to replication stress-inducing drugs in PDAC cells. **A**. Correlation plot of CDA expression and log_10_IC_50_ Irinotecan in PDAC cell lines from CCLE. **B**. Cell confluence (%) of PDAC015T transfected with control siRNA or CDA siRNA, 72 hours after transfection. **: p<0.01, ****: p<0.0001, unpaired t-test. **C**. Representative captions of PDAC051T cells treated by siRNA targeting CDA and 10μM oxaliplatin. **D**. Individual tumor growth of tumors, 10 days following treatment by CDA siRNA and oxaliplatin (n=7) or receiving control siRNA and oxaliplatin (n=6).

